# Glycolipid MPIase is essential for the TAT (Twin-Arginine Translocation) pathway

**DOI:** 10.64898/2026.06.05.730299

**Authors:** Hanako Nishikawa, Nachi Yamamoto, Hayate Kunezaki, Yui Shirogane, Kotoka Kanno, Katsuhiro Sawasato, Miwa Yamada, Ken-ichi Nishiyama

**Affiliations:** Department of Biological Chemistry and Food Sciences, Faculty of Agriculture, Iwate University, Morioka, Iwate 020-8550, Japan

## Abstract

TAT (Twin-Arginine Translocation) is a preprotein translocation system dedicated to membrane translocation of prefolded proteins in plants and bacteria. The TAT translocon, consisting of the TatABC subunits, drives translocation using proton motive force. However, there have been no reports on the successful reconstitution of the TAT system. In this report, we show that MPIase, a glycolipid that has known to catalyze membrane protein integration, is essential for the TAT system. Our findings in recombinant *Escherichia coli* demonstrate that overproducing TatABC increases MPIase levels and that depleting MPIase results in TAT precursor accumulation in the cytosol. Furthermore, co-reconstitution of MPIase with TatABC revealed the translocation activities of TAT substrates in a proton motive force-dependent manner. This is the first successful reconstitution of the TAT system and will be advantageous for understanding its mechanisms.

## Introduction

The TAT (Twin-Arginine Translocation) system, found in plant chloroplasts, bacteria, and mitochondria of some protozoa and plants, is dedicated to the membrane translocation of presecretory proteins with a TAT signal sequence ^1–4^. This signal sequence is composed of three regions, *n*, *h* and *c* regions from N-terminus: The *n* region contains the consensus motif of “SRRxFLK”, which includes the consecutive twin arginine residues that are essential for TAT translocation ^5^. The central *h* region (hydrophobic region) of the TAT signal sequence is in average less hydrophobic than that of the Sec signal sequence ^6^. The *c* region of the TAT signal sequence often contains basic amino acids that have been shown to function as a Sec-avoidance motif ^7^, preventing recognition of the TAT substrates by the Sec pathway ^8^. This region contains a cleavage site by Lep. Unlike the presecretory proteins that are translocated through the SecYEG or Sec61 translocon, TAT substrates maintain their folded structures during translocation. In many cases, TAT substrates fold and are transported with specific cofactors ^5,9^. It is known that *E. coli* K12 possess 27 TAT substrates ^9^. TAT substrates are translocated through the translocation machinery formed by TatABC using the proton motive force (PMF) as the driving force ^2,10–12^. The TAT machinery consists of the TatA and TatC families, all of which are membrane proteins ^13^. In *Escherichia coli*, the TatA family includes TatA and TatB, as well as TatE ^14–16^. TatE is a functional homologue of TatA that is not essential for TAT translocation ^14,17^. TatA and TatB are structurally similar but functionally distinct, since both are essential for the TAT translocation ^18–20^. On the other hand, TatA and TatE have overlapping functions in the TAT system ^14^. All three proteins are integral membrane proteins containing a single transmembrane helix (TMH), followed by a cytoplasmic amphipathic helix (APH) ^21,22^. TatC is a polytopic membrane protein with six TMHs ^23,24^. TatC forms a complex with TatB at an equal molar ratio, which recognizes the signal peptides of TAT substrates ^25–30^. In contrast, TatA forms a homomultimer that associates with the TatBC complex with different stoichiometries depending on the size of the substrate ^22,31–36^. It is also reported that that a signal peptide alone can induce TatA multimerization ^31^. Binding of substrates to the TatBC complex causes TatA to assemble around the complex in a PMF-dependent manner ^28,37^. The TatA cluster then forms a translocon through which the TAT substrates can pass when bound to the TatBC complex ^28,37–40^. According to MD simulations, the short TMHs and APHs of the TatA cluster have been shown to thin the membranes, thereby destabilizing the lipid bilayer of the cell membranes and allowing the translocation of prefolded proteins ^41,42^. Once translocation is complete, the substrate is released from the TatABC complex, and the signal sequence is cleaved by membrane-located leader peptidase ^10,43^. Assembly of the TAT machinery is a transient process that occurs exclusively during substrate translocation ^37,44^. As mentioned above, extensive studies performed by numerous researchers have significantly advanced our understanding of the molecular mechanisms underlying of the TAT system, particularly following the identification of a Tat component in 1997 ^45^. However, a reconstitution system for the TAT pathway has yet to be reported. Therefore, we hypothesized that a missing piece is necessary for successfully reconstituting the TAT system.

MPIase (*M*embrane *P*rotein *I*ntegr*ase*) is a glycolipid that catalyzes the integration of membrane proteins ^46^. Its glycan chain consists of approximately ten repeats of three *N*-acetylated amino sugars and is linked to diacylglycerol (DAG) by a pyrophosphate linker ^46^ (Fig. S1). This glycan chain has a molecular chaperon-like function, i.e., the glycan interacts directly with the substrate membrane proteins prior to insertion to prevent aggregation ^46–48^. The glycan is flexible on the membrane surface ^49^. DAG in the membrane bilayer blocks the disordered, spontaneous insertion by reducing the mobility of the acyl chain within the bilayers, while MPIase restores the mobility, allowing insertion with the substrate binding ^49^. In combination with the pyrophosphate linker, MPIase perturbs the membrane structure, enabling membrane proteins to insert into membranes. The negatively charged pyrophosphate of MPIase also attracts the positive charges near the TMH of the substrates ^47,48^.

MPIase can drive the membrane insertion of simple membrane proteins with a single TMH. MPIase also acts cooperatively with proteinaceous factors responsible for membrane insertion, such as the SecYEG translocase and the YidC insertase, in both Sec-dependent and - independent membrane insertion ^50,51^. MPIase is co-purified with SecYEG ^52^ and YidC ^53^. , indicating that MPIase interacts with these proteins directly. MPIase also stimulates protein translocation by affecting the dimer structure of SecYEG ^52,54,55^. Thus, MPIase is generally involved in both membrane integration and translocation ^56,57^, leading us to hypothesize that MPIase is also involved in the TAT pathway.

## Results

### MPIase is involved in the TAT pathway *in vivo*

Recently, we found that CdsA and its paralogue YnbB, CDP-DAG synthases, are involved in MPIase biosynthesis and constructed the strain KS23 (Δ*cdsA*::*cat* Δ*ynbB*) in which both the *cdsA* and *ynbB* genes were disrupted ^58^. The expression level of MPIase in this strain can be controlled by the addition of arabinose, since the *cdsA* gene is placed under the control of the *araBAD* promoter on a low-copy plasmid (pAra-CdsA). When arabinose is removed from the culture, MPIase becomes depleted (’ΔMPIase’ conditions) ^58^. We investigated the effect of MPIase depletion on the TAT pathway by using KS23/pAra-CdsA and its parent strain (EK413) (Fig. 1). When the TAT substrate SufI was expressed in EK413, the signal sequence-cleaved mature form of the protein was observed, in addition to the precursor. The level of the mature form increased when TatABC was provided from the *tet* promoter on a plasmid (pTet-TatABC) in addition to chromosomal expression. This finding is consistent with a previous report ^59^. The amount of mature SufI did not change upon SufI induction, indicating that the level was saturated. A large part of untranslocated precursors was aggregated (Fig. S2). In marked contrast, no mature form was observed upon MPIase depletion in KS23/pAra-CdsA, even when TatABC was expressed from pTet-TatABC (Fig. 1A, top). Upon SufI induction, a smaller band marked by a star appeared in addition to the precursor; however, this band is likely a degradation product of the precursor that had not been translocated, as it is larger than the mature band. The addition of arabinose restored MPIase expression (Fig. 1A, middle left) and generated the mature form of SufI (Fig. 1A, top). These results strongly suggest that MPIase is involved in TAT translocation. Similar results were obtained, when using TorA, another TAT substrate: no mature form was observed upon MPIase depletion (Fig. 1A, bottom left). The low level of TAT substrates under the MPIase depletion is due to the fact that MPIase-depleted cells hardly proliferate. Although TorA translocation appears to be less efficient than SufI translocation, this might be because TorA’s folding efficiency is lower in the absence of trimethylamine N-oxide (TMAO), as previously reported ^60^. TatABC, which had been expressed prior to arabinose removal, was still present after MPIase depletion (see Fig. 2C). To confirm that the processing to the mature form represents the TAT translocation, we examined the TAT signal mutants SufI (KK) (Fig. 1A, middle right) and TorA (KK) (Fig. 1A, bottom right). In these mutants, the twin arginine residues in the signal sequence were replaced with twin lysine residues, resulting in an inability to translocate across the IM. We observed that these KK mutants did not convert to the mature form, even in TatABC-overproducing strains (Fig. 1A, middle). These results indicate that the TAT translocation was properly studied. We investigated the involvement of MPIase in more detail using pulse-chase experiments (Fig. 1B). To facilitate TAT translocation, TatABC was overexpressed from the *tac* promoter on a plasmid (pTac-TatABC). In the wild-type strain harboring pTac-TatABC (Fig. 1B, top left), the precursor form of SufI was efficiently converted to the mature form. No mature form was observed when the signal mutant SufI (KK) was used (Fig. 1B, top right), confirming that the TAT translocation was analyzed correctly. When MPIase was depleted, no mature form was observed even after a 10-min chase (Fig. 1B, bottom, ‘ΔMPIase’), confirming MPIase’s involvement in the TAT system. When MPIase synthesis was induced (Fig. 1B, bottom, ‘+MPIase’), the mature form appeared, but the conversion rate was much slower than under wild-type conditions. The reason for this will be discussed later. To investigate the role of MPIase in the TAT system, we performed cell fractionation into periplasmic and spheroplast fractions (Fig. 1C). When the MPIase-depleted strain expressing TatABC and SufI was fractionated, the precursor was recovered in the spheroplast fraction, and no mature form was observed. This indicates that SufI was not translocated (Fig. 1C, left, ‘ΔMPIase’). Conversely, mature SufI was translocated to the periplasm, when MPIase was expressed (Fig. 1C, left, ‘+MPIase’). Successful fractionation was confirmed by detecting the periplasmic protein β-lactamase and the cytoplasmic protein SecB. Similar results were obtained when TorA-GFP, in which the TorA signal is connected to GFP, was examined (Fig. 1C, right), confirming that TorA-GFP is not translocated upon MPIase depletion. These cells expressing TorA-GFP were also used to detect the subcellular localization directly by monitoring its fluorescence (Fig. 1D). In the presence of MPIase, the envelope fraction exhibited green fluorescence, indicating that GFP is localized in the periplasm or on the inner membrane at least (left). Conversely, in the absence of MPIase, the entire cell exhibited uniform fluorescence, indicating that TorA-GFP is localized in the cytoplasm under these conditions (right). These results are consistent with those of the subcellular fractionation (see Fig. 1C). Taken together, we conclude that MPIase is involved in the TAT system *in vivo*.

**Fig. 1.**
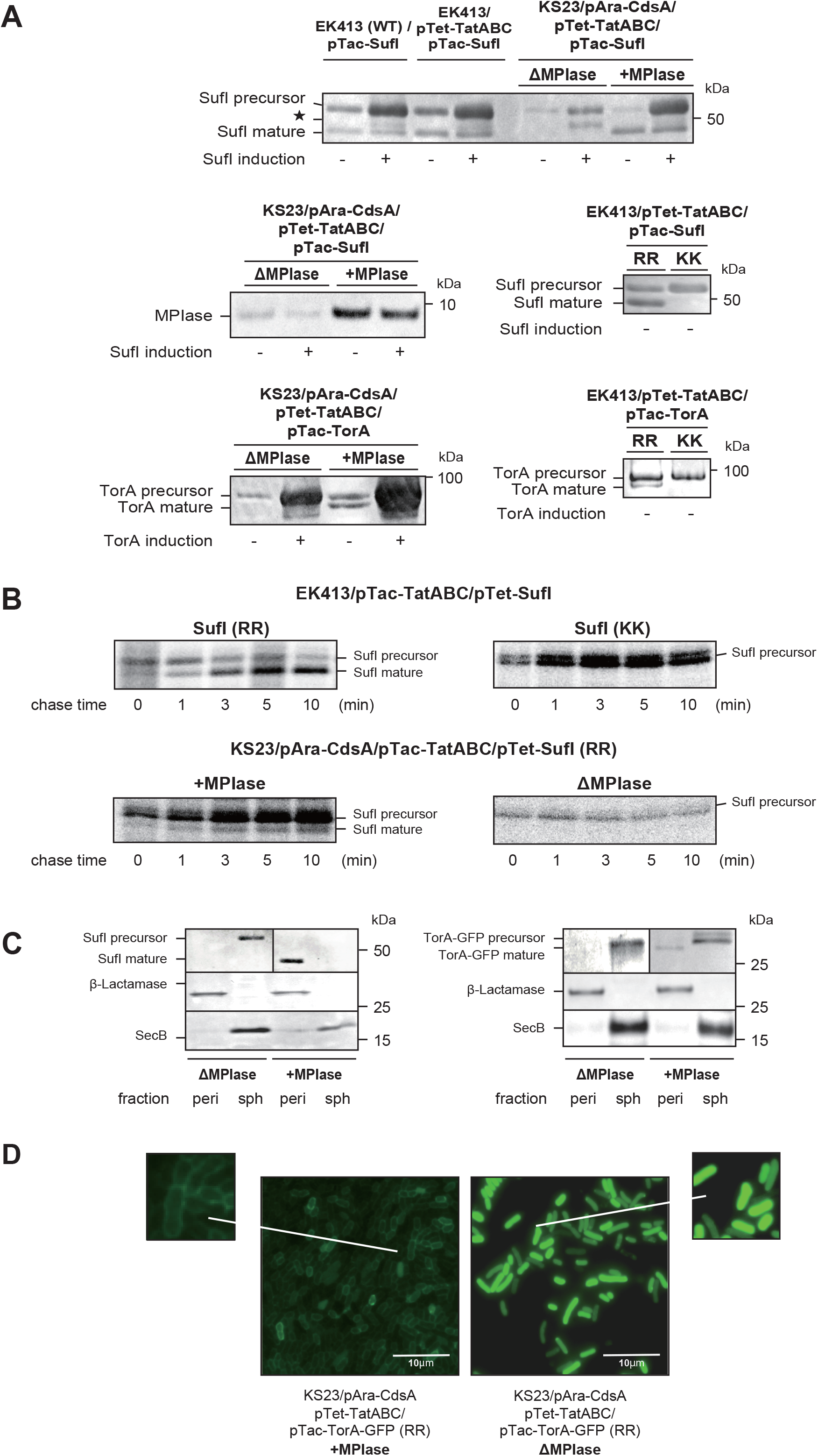
MPIase depletion causes the accumulation of TAT substrate precursors. **(A)** Detection of SufI and TorA precursors upon MPIase depletion. SufI (top) and TorA (bottom) were expressed in the indicated strains. IPTG (1 mM) was added at early log phase, and cultivation was continued for 2 h. The cells were subjected to TCA precipitation, followed by SDS-PAGE/immunoblotting. The precursor and mature forms were then detected as indicated. MPIase depletion was confirmed in KS23 (middle left). Defects in the processing of SufI (middle right) and TorA (bottom right) with the KK signal were also confirmed. The symbol ’ ’ denotes degradation products of the SufI precursor. **(B)** Pulse-chase experiments of SufI processing after MPIase depletion. EK413/pTac-TatABC harboring pTet-SufI(RR) (top left) or pTet-SufI(KK) (top right), and KS23/pAra-CdsA/pTac-TatABC harboring pTet-SufI(RR) (+MPIase; bottom left, ΔMPIase; bottom right) were pulse-labeled for 30 s and then chased for the indicated periods. SufI(RR) and SufI(KK) were isolated using the attached His tags, followed by SDS-PAGE/autoradiography. **(C)** Subcellular localization of SufI and TorA-GFP after MPIase depletion. KS23/pTet-TatABC/pTac-SufI(RR) (left) and KS23/pTet-TatABC/pTac-TorA-GFP(RR) (right) were cultivated in the presence (+MPIase) or absence (ΔMPIase) of arabinose. The TAT substrates were leaky expressed from the *tac* promoter. The cells were then fractionated into periplasmic (peri) and spheroplast (sph) fractions. The indicated proteins were detected by immunoblotting. **(D)** Subcellular localization of TorA-GFP after MPIase depletion as revealed by fluorescence microscopy. The indicated strains were cultivated, TorA-GFP was induced, and then it was observed by fluorescence microscopy. The scale bars represent 10 μm. A magnified view of part of the ’+MPIase’ sample is shown on the left.

**Fig. 2.**
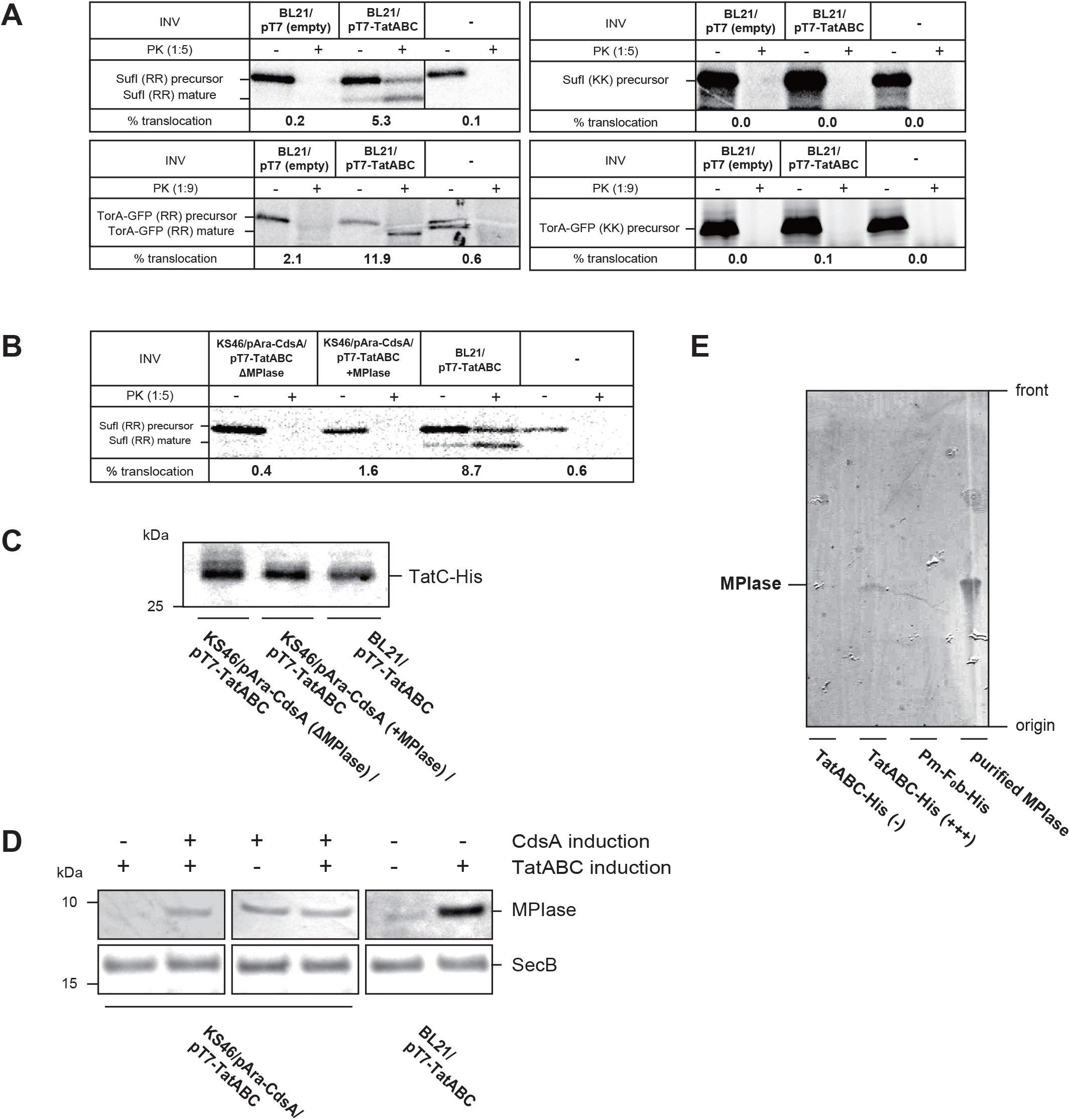
TAT substrates are translocated into INV *in vitro* when TatABC and MPIase are both overproduced. **(A)** Translocation of SufI and TorA-GFP into TatABC-overproducing INV. SufI (top) and TorA-GFP (bottom) were synthesized *in vitro* in the presence of INV prepared from the indicated strains. INV was omitted in the ’-’ samples. The signal sequences on the left are wild-type (RR), while the sequences on the right are mutants (KK). Translocated materials were generated after PK digestion (+). One-fifth of the reaction mixture was analyzed as the translation standard (-). Translocation activities, expressed as a percentage of the amount of the synthesized proteins, are shown at the bottom of each autoradiogram. **(B)** Depletion of MPIase resulted in the failure of TAT translocation. INV were prepared from KS46/pAra-CdsA harboring pT7-TatABC, which was cultivated in the presence (+MPIase) or absence (ΔMPIase) of arabinose. After washing arabinose, IPTG was added at 1 mM to induce TatABC, and cultivation was continued for 2 h prior to INV preparation. SufI translocation was then performed as described in (A). **(C)** The level of TatC in INV used in (A) and (B). Each INV (10 μg) was analyzed by SDS-PAGE, followed by immunoblotting using anti-RpmJ-His antibodies. The position of TatC-His was shown. **(D)** MPIase is upregulated upon TatABC overproduction. The expression levels of MPIase in the strains used in (B) were determined by immunoblotting. CdsA and TatABC were induced as indicated. SecB levels were determined as a loading control. **(E)** MPIase is co-purified with TatABC. INV prepared from BL21/pT7-TatABC-His induced with 1 mM IPTG (+++) or not induced (-), and BL21/pT7-Pm-F_0_b-His induced with 1 mM IPTG, solubilized with DDM, were applied to a TALON column, then eluted. The eluates were analyzed by TLC, followed by immunodetection of MPIase. The positions of the origin/front and MPIase are shown. Purified MPIase was also analyzed at right.

### Upregulation of MPIase upon TatABC overproduction is essential for *in vitro* TAT translocation

The TAT pathway can be reproduced *in vitro* using inverted membrane vesicles (INV), as depicted in Fig. S3. However, it has been shown that overproduction of TatABC is necessary for translocation into INV ^10,59^. To reproduce these observations, we prepared INV from wild-type strains with an empty vector (Fig. 2A BL21/pT7 (empty)) and from TatABC-overproducing strains (Fig. 2A, BL21/pT7-TatABC), which were then subjected to the SufI translocation assay *in vitro*. TatABC overproduction with this vector was confirmed by immunoblotting (Fig. S4).

As reported ^10,59^, SufI translocation into INV was observed only when TatABC was overproduced (Fig. 2A, top left). In the absence of INV, SufI was completely digested by PK (-). These results indicate that SufI is translocated in a TatABC-dependent manner. SufI (KK) was not translocated, even when TatABC-overproducing INV were used (Fig. 2A, top right), indicating that the TAT pathway is adequately analyzed in our experimental system. We observed the same trend with another TAT substrate, TorA-GFP (Fig. 2A, bottom). To further test the involvement of MPIase in the TAT pathway in this *in vitro* system, we used a *cdsA*/*ynbB* knockout strain with a BL21 (DE3) background (KS46) harboring pAra-CdsA. We then prepared MPIase-depleted INV with TatABC overproduction, followed by SufI translocation (Fig. 2B).

Since TatABC was overproduced prior to MPIase depletion, these INV contain the overproduced amount of TatABC (Fig. 2C) but not MPIase (Fig. 2D). We observed that SufI was not translocated into these MPIase-depleted INV, which further supports the involvement of MPIase. Consistently, the anti-MPIase antibody inhibited SufI translocation (Fig. S5). However, we also found that the translocation activity with MPIase-induced INV (Fig. 2B, ‘+MPIase’) did not recover to the level observed with the wild-type INV with TatABC overproduction (Fig. 2B, ‘BL21/pT7-TatABC’). The reason for the incomplete recovery was that TatABC overproduction caused MPIase upregulation (Fig. 2D). Significant MPIase upregulation occurred when TatABC overproduction was induced in BL21(DE3). However, MPIase expression remained unchanged with TatABC overproduction in KS46, in which *cdsA* was induced from the plasmid pAra-CdsA (Fig. 2D). Therefore, authentic promoters for *cdsA* should be responsible for this upregulation, because CdsA is a rate-limiting enzyme for MPIase biosynthesis ^58^. Since the levels of Tat components in BL21(DE3) and KS46/pAra-CdsA grown with arabinose were similar (Fig. 2C), it is reasonable to conclude that MPIase upregulation is necessary for the translocation activity *in vitro* of the TAT pathway into INV. This explains the low activity of MPIase-induced INV (Fig. 2B, ‘KS46/pAra-CdsA +MPIase’). These results are consistent with the fact that the SufI translocation was not fully restored in the pulse-chase experiments when MPIase was induced from pAra-CdsA (see Fig. 1B). MPIase upregulation was specific to TatABC overproduction; MPIase levels were barely affected by SecYEG overproduction ^52^ (Fig. S6). These results strongly suggest that TatABC and MPIase function cooperatively. To examine whether the interaction between TatABC and MPIase is direct and physical, we detected MPIase in the TatABC-His preparation purified via metal affinity chromatography. As shown in Fig. 2E, MPIase was co-purified with TatABC. Furthermore, the amount of MPIase precipitated with TatABC increased significantly with TatABC overproduction (Fig. 2E). When His-tagged F_0_b from *Propionigenium modestum* (Pm-F_0_b-His) was expressed and used as a control, no detectable amount of MPIase was precipitated. Under these conditions, both TatABC-His and Pm-F_0_b-His were purified to homogeneity (Fig. S7). These results indicate that TatABC and MPIase directly and specifically interact with each other. Based on these results, we concluded that the TatABC requires MPIase to function.

### MPIase is the missing piece for successfully reconstituting the TAT pathway

Results obtained thus far, both *in vivo* and *in vitro*, demonstrate the essentiality of both TatABC and MPIase. Therefore, we assumed that the TAT reconstitution might be possible if TatABC and MPIase were both present and PMF was imposed. To test if our reconstitution system would work, we started with a crude system using proteoliposomes reconstituted from the solubilized membranes. First, we prepared proteoliposomes from solubilized membranes with overproduced amounts of TatABC and purified F_0_F_1_-ATPase from thermophilic bacteria (see Fig. 3B, left), followed by inclusion of DAG for stable generation of PMF ^61^. However, these proteoliposomes were inactive with regard to SufI translocation (Fig. 3A, top left, ‘+PL/DAG’).

**Fig. 3.**
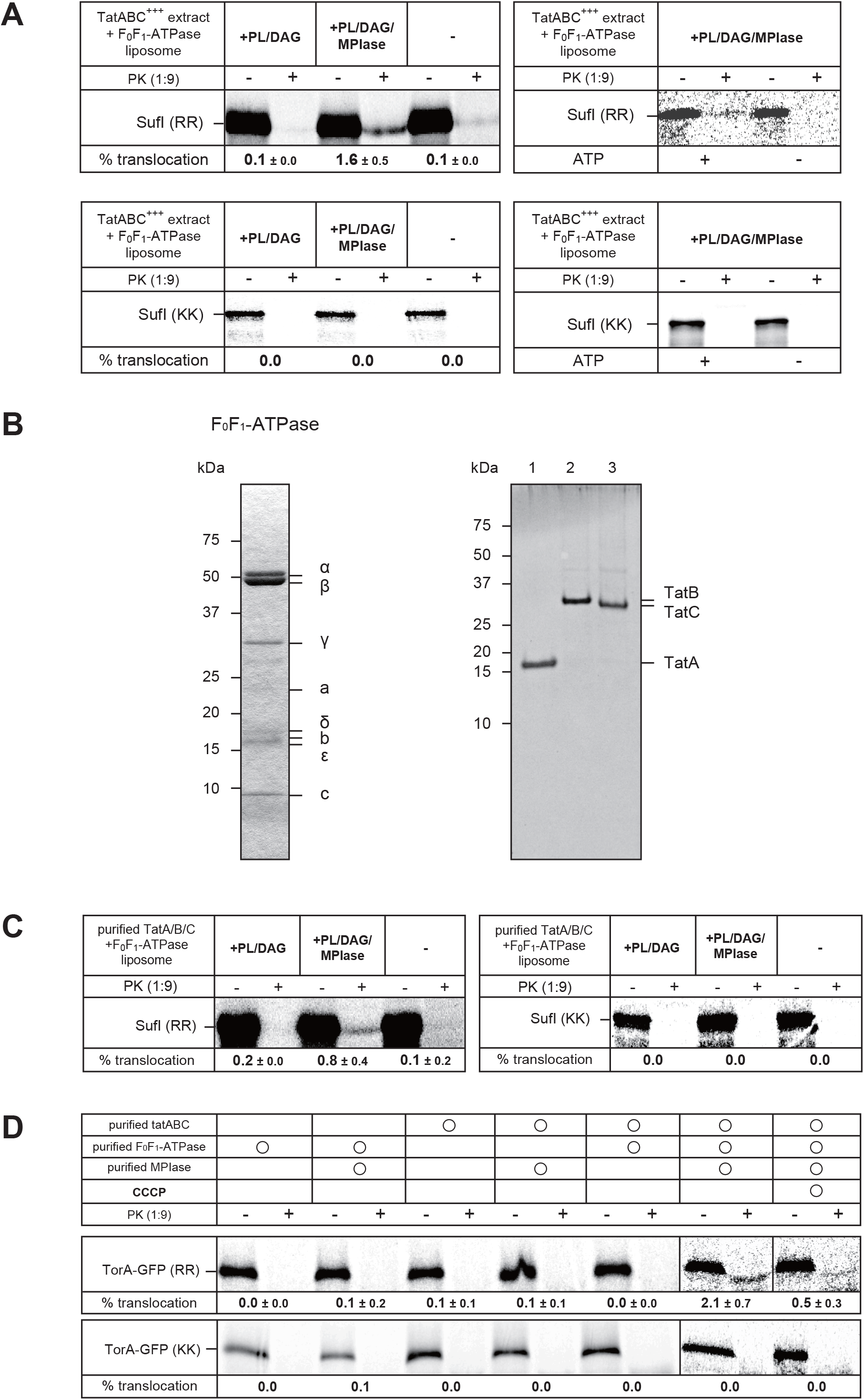
Reconstitution of the TAT system. **(A)** Membrane fusion of TatABC proteoliposomes with MPIase liposomes enabled the detection of the SufI translocation. INV, prepared from BL21/pT7-TatABC, were solubilized with OG, and then mixed with purified F_0_F_1_-ATPase. The proteoliposomes were then formed by removing the detergent. PL/DAG liposomes or PL/DAG/MPIase liposomes were fused to the reconstituted proteoliposomes. SufI translocation was performed co-translationally (left) or post-translationally (right) in the presence of these proteoliposomes. SufI(RR) (top left) or SufI(KK) (bottom left) was used as the substrate. One-ninth of the reaction mixture was analyzed as the translation standard (-) for the co-translational reactions. For the post-translational reactions, ATP was added as indicated. Translocation activities are shown at the bottom of the left panels. **(B)** Purification of F_0_F_1_-ATPase and the TAT components. F_0_F_1_-ATPase from a thermophilic Bacillus PS3 (left), and *E. coli* TatA-His (lane 1), TatB-His (lane 2), and TatC-His (lane 3) (right) were subjected to SDS-PAGE, followed by Coomassie Brilliant Blue staining. **(C, D)** Complete reconstitution of the TAT system using the purified components. **(C)** Purified TatA, TatB, and TatC (in a 1:1:1 ratio) and F_0_F_1_-ATPase were mixed with phospholipids. Then, detergent was removed to form proteoliposomes. The reconstituted proteoliposomes were then fused with PL/DAG liposomes or PL/DAG/MPIase liposomes. Translocation activity was assayed with SufI(RR) (left) or SufI(KK) (right). **(D)** Proteoliposomes were reconstituted with the indicated components, followed by TorA-GFP translocation. The TatA:TatB:TatC ratio was 10:1:1. CCCP (20 μM), a protonophore, was added to the sample on the left. TorA-GFP(RR) (top) or TorA-GFP(KK) (bottom) was used as the substrate. Translocation activities are shown at the bottom of each gel. Errors denote +/- SD (N=3).

We found that fusing MPIase-containing liposomes to the proteoliposomes greatly enhanced SufI translocation (Fig. 3A, top left, ‘+PL/DAG/MPIase’). This may be because MPIase was not efficiently recovered in the proteoliposome fraction due to the hydrophilic nature of MPIase’s glycan moiety ^46^. These results again indicate that MPIase is essential for the TAT system. Here, we could not observe the signal sequence processing. This may be due to the dilution of membrane proteins, including Lep, in proteoliposome membranes since signal cleavage and translocation are not coupled ^39^. Even when MPIase-containing liposomes were fused, SufI (KK) was not translocated into the proteoliposomes (Fig. 3A, bottom left). We also examined the post-translational translocation using a partially purified SufI precursor (Fig. 3A, top right). We observed SufI (RR) translocation into TatABC/MPIase proteoliposomes in an ATP-dependent manner (i.e., a PMF-dependent manner), but not SufI (KK) translocation. Thus, the TAT reconstitution was achieved through a combination of TatABC, MPIase and F_0_F_1_-ATPase.

Next, we purified TatA, TatB and TatC individually through an attached His tag (Fig. 3B, right). We confirmed that expressing each His-tagged TatA, TatB, and TatC complemented the SDS-sensitive phenotype of the corresponding knockout strain (Fig. S8). Using these purified proteins, we reconstituted proteoliposomes containing TatABC and F_0_F_1_-ATPase. Then, we fused them with MPIase/DAG liposomes. The TatABC-MPIase proteoliposomes containing F_0_F_1_-ATPase exhibited activity with respect to SufI translocation (Fig. 3C, left). This activity was comparable to that of proteoliposomes reconstituted with solubilized membranes (Fig. 3A). On the other hand, SufI (KK) was not translocated into the proteoliposomes (right). In these experiments, reconstituted proteoliposomes were prepared using a TatA/B/C weight ratio of 1/1/1. To obtain higher activity, we examined various TatA/B/C ratios. We found that increasing the amount of TatA gave slightly higher activity. We then prepared proteoliposomes containing TatA/B/C at a ratio of 10/1/1. Similar results with improved activity were obtained when using TorA-GFP (Fig. 3D). Proteoliposomes containing both TatABC and MPIase allowed for the translocation of TorA-GFP (RR) but not TorA-GFP (KK). The translocation of TorA-GFP (RR) was severely inhibited by CCCP (carbonyl cyanide 3-chlorophenylhydrazone), a protonophore, even if the reaction mixture contains DTT that inactivates CCCP, indicating that the translocation is dependent on PMF (right). Furthermore, omitting MPIase, TatABC or F_0_F_1_-ATPase resulted in a loss of translocation activity (left). These results indicate that TatABC, MPIase and PMF are necessary and sufficient for TAT translocation.

### MPIase serves as the TAT signal receptor

Although the participation of MPIase in the TAT pathway has been demonstrated both *in vivo* and *in vitro*, the mechanisms by which MPIase contributes to TAT translocation remains unclear. Our previous study showed that the glycan chain and the pyrophosphate residue of MPIase captures the substrate membrane protein on the membrane surface before membrane insertion ^48,56^. Although the TatBC complex has been shown to act as the TAT signal sequence receptor ^26,30^, we hypothesized that MPIase could also be involved in the targeting of the TAT substrate proteins to the membrane. Thus, we examined whether or not MPIase is involved in the membrane localization of TorA-GFP. Cells expressing TorA-GFP were fractionated into soluble (cytosol plus periplasm) and membrane fractions. Then, we measured the GFP fluorescence. We confirmed that the cell debris and microaggregates could be removed by low-speed centrifugation before membrane sedimentation (Fig. S2). When we induced MPIase in KS23 cells by adding arabinose, ∼30% of the fluorescence was recovered in the membrane fraction (Fig. 4A, ‘RR, +MPIase’), similarly to the ‘EK413’ case. Note that the MPIase level was insufficient under these conditions, resulting in very low translocation efficiency (see Fig. 1B). Conversely, when MPIase was depleted, only <10% of the fluorescence was recovered in the membrane fraction (Fig. 4A, ‘RR, ΔMPIase’). Similar results were obtained using TorA-GFP with a mutated signal sequence (KK) (Fig. 4A). These results strongly suggest that MPIase serves as a receptor for the TAT signal, but that MPIase does not discriminate the functional signal. Similar to the ‘+MPIase’ and ‘EK413’ cases, ∼30% of the fluorescence was recovered in the membrane fraction when the *tatC* knockout mutant was used (Fig. 4A, ‘ΔTatC’). These results suggest that MPIase recognizes the TAT signal before TatC does. When GFP without the TAT signal was expressed, >90% of fluorescence was recovered in the supernatant fraction, regardless of MPIase levels and TatABC overproduction (Fig. S9), indicating that the membrane localization depends solely on the TorA signal.

**Fig. 4.**
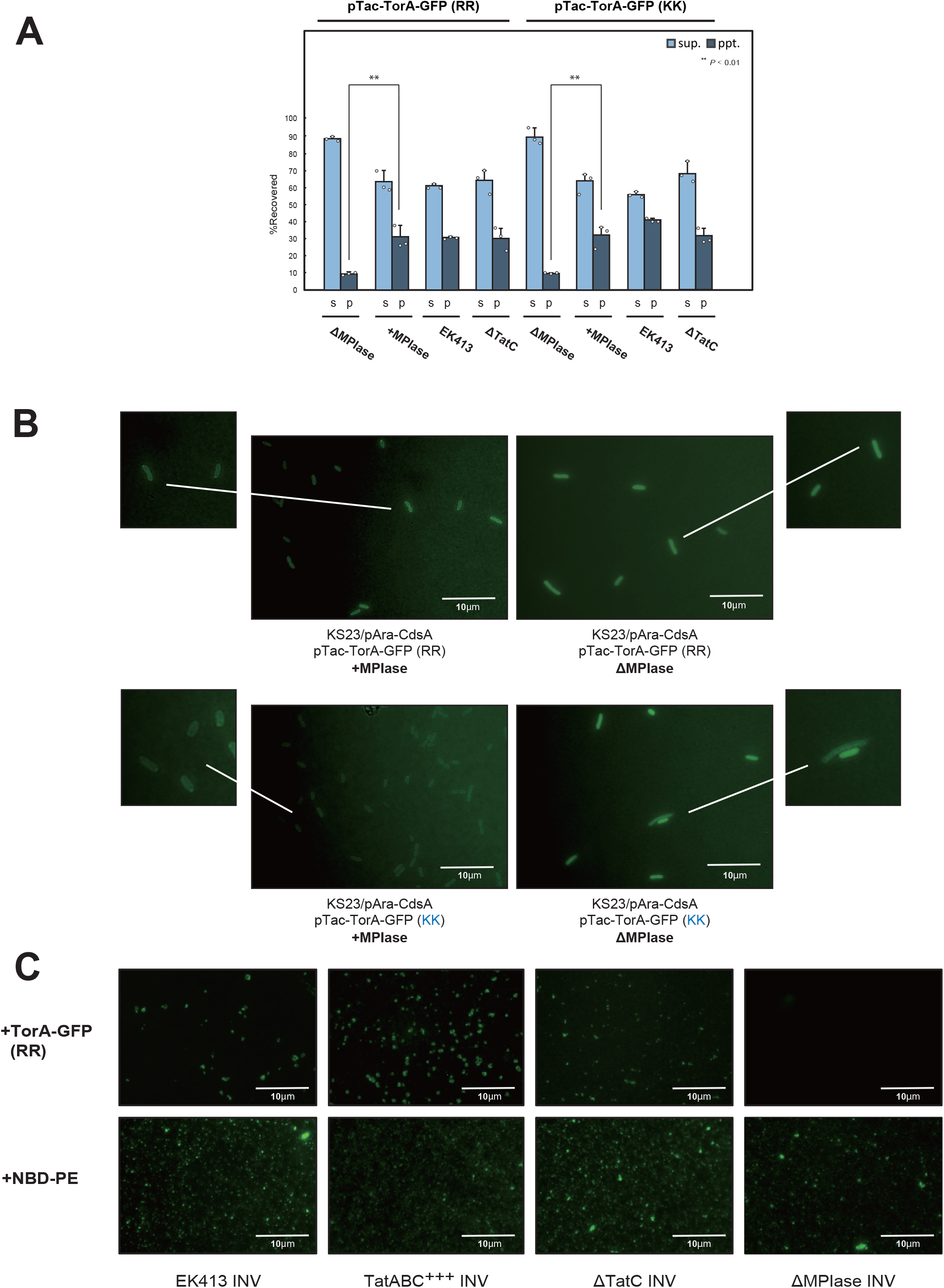
MPIase is a membrane receptor for TAT substrates. **(A,B)** TorA-GFP is targeted to the membrane in an MPIase-dependent manner. TorA-GFP(RR) (A; left half) or TorA-GFP(KK) (A; right half) was expressed in the specified strains. The cells were then disrupted to obtain the supernatant (sup; cytosol and periplasm) and the precipitate (ppt; membranes) fractions. The fluorescence intensity of each fraction was measured. The ratio to the whole cell was determined and shown as percentages. The means and standard deviations were calculated from at least three independent experiments. **, *P*<0.01. In (B), the indicated strains were cultivated, TorA-GFP was induced, and then it was observed by fluorescence microscopy. The scale bars represent 10 μm. Part of each picture was magnified on each side. Scale bars represent 10 μm. **(C)** TorA-GFP interacts with INV expressing MPIase. Purified TorA-GFP was mixed with the indicated INV, followed by fluorescence detection (top). As a control, each INV was mixed with NBD-PE, a fluorescent phospholipid, and observed (bottom). Scale bars represent 10 μm.

The MPIase-dependent membrane targeting was also observed using fluorescent microscopy (Fig. 4B). When MPIase was induced by adding arabinose, an envelope-shaped fluorescence signal was detected, regardless of whether it was wild-type or mutant (Fig. 4B, left), indicating that TorA-GFP (RR) and TorA-GFP (KK) were both targeted to membranes.

Conversely, under the MPIase-depleted conditions, whole-cell fluorescence was detected (Fig. 4B, right), indicating that both TorA-GFP (RR) and TorA-GFP (KK) located in the cytosol.

MPIase-dependent targeting was confirmed *in vitro* (Fig. 4C, top). When the purified TorA-GFP (Fig. S10) was mixed with wild-type INV (Fig. 4C, ‘EK413’), TatABC-overproducing INV (Fig. 4C, ‘TatABC^+++^’) and TatC-deficient INV (Fig. 4C, ‘ΔTatC’), in all of which contain MPIase, fluorescent dots were observed, indicating that TorA-GFP directly interacts with these INV. The number of dots increased with TatABC overproduction, which is consistent with MPIase upregulation induced by TatABC overproduction. In contrast, no such fluorescent dots were observed in the presence of MPIase-depleted INV (Fig. 4C, ‘ΔMPIase’), confirming the MPIase-dependent targeting of TorA-GFP. The presence of similar amounts of membrane vesicles was visualized by adding fluorescent phospholipids, NBD-PE (Fig. 4C, bottom).

## Discussion

In this study, we demonstrated that glycolipid MPIase is essential for the TAT system. All of the *in vivo*, *in vitro* and reconstitution studies supported this conclusion. Therefore, the previous unsuccessful attempts at TAT reconstitution were likely due to an insufficient amount of MPIase in TatABC proteoliposomes. Even when using solubilized membranes containing TatABC and MPIase to prepare proteoliposomes, MPIase recovery would be too low to detect the TAT activity.

MPIase is a glycolipid that catalyzes the membrane insertion of proteins with a single TM ^50,53^ as well as those with multiple TM domains ^51,55,57^. Therefore, depleting MPIase should impair the membrane insertion of all the TAT components, TatA, TatB and TatC, leading to the failure of TAT translocation. However, in our experimental system, TatABC was still present after MPIase depletion (see Fig. 2C), which supports the direct involvement of MPIase in the TAT translocation. Previous studies have reported that the Sec translocation slows down but does not stop completely, when MPIase is depleted ^58^, suggesting that the Sec machinery, including SecYEG, remains present during the initial stages of MPIase depletion, as demonstrated in this study. The results of the reconstitution study, in which TatABC was reconstituted in proteoliposome membranes, rule out the indirect involvement of MPIase in TAT translocation. In this system, the TAT translocation occurred in TatABC-, PMF- and MPIase-dependent manners.

How is then MPIase involved in the TAT system? Since MPIase has been identified as a factor that catalyzes membrane protein integration ^46,50^, it is possible that MPIase drives the insertion of the TAT signal sequences into membranes, thereby initiating the translocation reaction (Fig. 5, (i)). Depleting MPIase caused TorA-GFP to accumulate in the cytosol. Thus, MPIase serves as a receptor for TAT precursors, and the TAT signal sequences are inserted into membranes. In this process, both hydrophobic interaction between the *h*-region and the acetyl residues of the MPIase glycan, and electrostatic interaction between the positive charges of the *n*-region and the pyrophosphate residue of MPIase would be involved, as observed in the interaction between substrate membrane proteins and MPIase ^47^. Then, they are transferred to TatC (ii), where the functional TAT signal is verified (iii). In addition to its roles as a membrane receptor and an insertase for the TAT signals, MPIase may facilitate structural changes in TatABC, allowing the complex to form the translocon dedicated to the substrate protein (iv). The TAT substrate is then translocated through the translocon, and the mature protein is released into the periplasm upon Lep-mediated cleavage of the signal sequence ^10,43^ (v). These processes require PMF, as previously reported ^62^. Thus, MPIase may be involved not only in the membrane targeting of TAT substrates, but also in forming the TatABC translocon for each substrate.

**Fig. 5.**
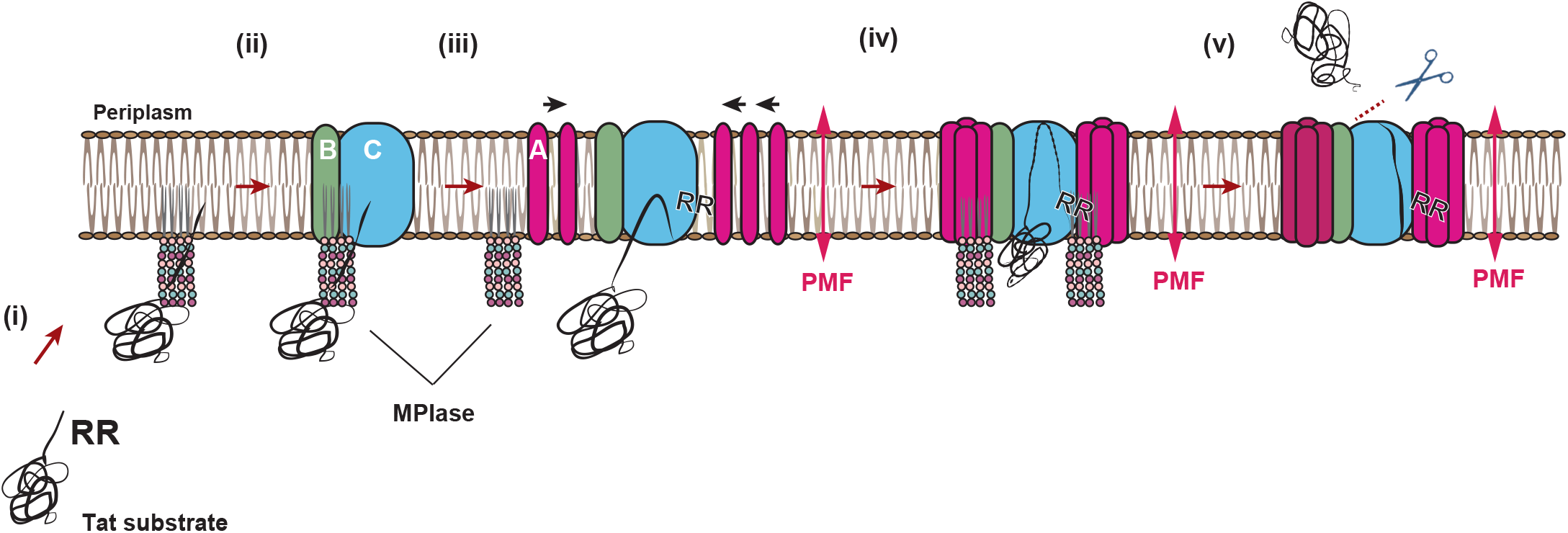
Working model of mechanisms underlying MPIase-dependent TAT translocation. MPIase recognizes the signal sequence of the TAT substrates and insert them into the membrane. The TatBC complex on the membrane then determines whether the signal sequence of the membrane-inserted TAT substrate is valid. The TatABC translocon then forms a pore the same size as the substrate. This allows the substrate protein to be translocated across the membrane. Finally, Lep cleaves the signal sequence, and the mature protein is released into the periplasm. See the text for details.

Membrane lipids PE may be important for these processes, since PE is essential for TAT translocation ^63^. In the reconstitution studies, the absence of DAG in (proteo)liposomes allowed for the spontaneous insertion of membrane proteins and signal sequences ^55,64^. Therefore, the failure of the TAT reconstitution strongly suggests that MPIase has functions beyond serving as a membrane receptor and an insertase for TAT translocation. Given that MPIase induces dynamic structural changes in the SecYEG translocon ^52,54^, it is likely that MPIase also modifies the structure of TatABC to form a tailor-made translocon. This possibility is consistent with the observations showing that TatABC overproduction induces MPIase upregulation and that MPIase co-precipitates with TatABC.

It is well known that the lipid environment of the membranes significantly affects the efficiency of TAT translocation. First, acidic phospholipids are important for TAT translocation because TAT signals interact with the acidic phospholipids ^65^. MPIase has a pyrophosphate residue and would therefore contribute to this interaction in a similar way. At the same time, the hydrophobic domain of the TAT signals is solubilized by the glycan chain, as occurs with membrane proteins ^47^. Conversely, a decrease in PE (i.e., an increase in acidic phospholipids, PG and CL) decreases efficiency ^66,67^, presumably due to an overly strong interactions between the TAT signals and the acidic phospholipids. Alternatively, PE may be important for TAT translocation, since it stabilizes several membrane proteins ^68^. Thus, PE may be involved in the expression of the TatABC function. In the thylakoid system, the reduced unsaturation in membrane lipids results in a 50% decrease in the efficiency of TAT translocation ^69^, suggesting that membrane fluidity is also important for TAT translocation. We have demonstrated that MPIase increases the membrane fluidity through the flexible movement of its long sugar chain ^49^. This property would benefit TAT translocation.

MPIase is upregulated in the cold. During this process, both *cdsA* and *ynbB* are induced at the transcriptional level. In the case of *cdsA* induction, two cold-inducible promoters located upstream of the *cdsA* gene are responsible ^70^. MucA, a small peptide that overlaps with *cdsA*, enhances MPIase synthesis by CdsA ^71^. When MPIase is upregulated by TatABC overproduction (see Fig. 2D), *cdsA* and/or *ynbB* are also induced, as evidenced by the diminished increase in the MPIase level in *cdsA*/*ynbB* knockouts even when CdsA expression is complemented from the *araBAD* promoter on a plasmid. In this case, the *cdsA* promoters other than identified cold-inducible ones would be responsible, together with MucA. Alternatively, other mechanisms could operate to upregulate MPIase.

We found that the TAT translocation could be reconstituted with TatABC and MPIase alone when PMF was imposed, indicating that these factors are sufficient for TAT translocation. Although the activity was comparable to that of INV and proteoliposomes with the crude extract, it was still low. Therefore, these results suggest the presence of factor(s) involved in TAT translocation that stimulate the reaction, as the soluble factors that solubilize the TAT substrates and target them to membranes remain unidentified.

The TAT system is widely used as a carrier system for secreting foreign proteins into the medium ^72–74^. Now that we know that MPIase is involved in the TAT system, it will be easier to engineer the TAT system by modulating the MPIase activity to increase the level of secreted proteins.

The TAT system was first identified in the thylakoid membranes of chloroplasts as a ΔpH-dependent pathway ^2,75^. The components of the TAT system in chloroplasts are highly homologous to TatABC in *E. coli* ^26,37^, and the translocation reaction is also driven by PMF ^2^. Thus, the mechanisms underlying the TAT systems in *E. coli* and chloroplasts likely have similar underlying mechanisms, strongly suggesting the presence of an MPIase homologue in chloroplasts. This similarity could facilitate the identification of an MPIase homologue in chloroplasts.

## Methods

### Materials and methods

The strains, plasmids, and oligonucleotides used in this study are listed in Tables S1, S2, and S3, respectively. INV were prepared as described ^59^. The plasmid pCspB-TorAss-GFP encoding TorA-GFP was obtained from Dr. Y. Kikuchi (Ajinomoto Co., Inc.). Bio Beads SM-2 were purchased from BioRad. [^35^S] EXPRE^35^S^35^S Protein Labeling Mix, a mixture of [^35^S] methionine and [^35^S] cysteine (∼37 TBq/mmol), was obtained from Revvity. Proteinase K (PK) was purchased from Roche Diagnostics. The cell-free protein synthesis kit “*E. coli* S30 Extract System for Circular DNA” was purchased from Promega Corporation. The *E. coli* polar lipid extract (phospholipids), 18:1 NBD-PE, and dioleoylglycerol were obtained from Avanti Polar Lipids, Inc. *n*-Octyl-β-D-glucopyranoside (OG) and *n*-dodecyl-β-D-maltopyranoside (DDM) were obtained from Dojindo Laboratories. IPTG was from Wako Pure Chemicals Industries, Ltd. TLC plates were from Merck. MPIase was purified from MC4100, as described ^46,50^. TorA-GFP, TatA, TatB, TatC, and the TatABC complex were purified from DH5α/pTac-TorA-GFP-His, DH5α/pTac-TatA-His, DH5α/pTac-TatB-His, DH5α/pTac-TatC-His, BL21/pT7-Pm-F_0_b-His, and BL21/pT7-TatABC-His, respectively. These proteins were induced with 1 mM IPTG, followed by INV preparation, solubilization and purification through the 6 x His-tag attached to C-termini on Talon columns. The conditions for INV solubilization and Talon column chromatography was carried out as described ^57^. Purification of F_0_F_1_-ATPase from a thermophilic bacterium PS3 was performed as described ^76^.

### Depletion of MPIase

Strains KS23/pAra-CdsA and KS46/pAra-CdsA were cultivated overnight at 37°C in LB medium containing 0.002% arabinose. After the cells were washed five times with fresh LB medium, they were inoculated into LB medium supplemented or not with 0.2% arabinose at a dilution of 1/1000 and then shaken at 37°C. To express the TAT substrates, IPTG (1 mM) was added after the cell culture was grown to OD_600 nm_ ∼0.5 and cultivated for another 1 h. An aliquot (500 µL) of each sample was treated with TCA to a final concentration of 5%. The TCA precipitate was recovered by centrifugation (10,000 x g, 4°C, 5 min), followed by washing with acetone. MPIase depletion was then confirmed by SDS-PAGE/immunoblotting, as described ^58^.

### Pulse-chase experiments

Cell cultures grown in LB medium to OD_600 nm_ ∼0.5 were harvested and resuspended in M9 minimal medium containing 18 amino acids (0.1 mM each) except for methionine and cysteine, and 0.1% yeast extract. After incubating for an additional 1 h at 37°C, the substrate proteins were labeled with [^35^S] methionine/cysteine (∼10 MBq/mL) for 30 s. Labeling was terminated by adding non-radioactive methionine and cysteine to a final concentration of 12 mM. At the specified chase time, an aliquot (500 µL) of the culture was withdrawn, and TCA was added to a final concentration of 10%. The TCA precipitates were washed successively with acetone and diethyl ether, and then suspended in 50 µL of 50 mM Tris-HCl (pH 7.5), 1% SDS, 1 mM EDTA (pH 8.0), and boiled for 3 min. The samples were then diluted with 1 mL of 50 mM HEPES-KOH (pH 7.5), 150 mM NaCl, 1% Triton X-100, 1 mM EDTA (pH 8.0), 1 mM phenylmethylsulfonyl fluoride, and were centrifuged at 16,000 x g for 5 min. Then, 10 µL of TALON resin was added to the supernatant, and the mixture was stirred at 4°C for 1 h. After centrifugation (3,000 x g, 4°C, 30 s), the TALON resin was washed successively with 500 µL of 50 mM Tris-HCl (pH 7.5), 150 mM NaCl, 1% Triton X-100. The washed resin was suspended in 20 µL of the sample buffer for SDS-PAGE. Radioactive bands, analyzed by SDS-PAGE, were detected by autoradiography using a phosphorimager (GE Healthcare), and quantified using ImageQuant (GE Healthcare).

### Preparation and fractionation of spheroplasts

After the cells were grown in LB medium to OD_600 nm_ ∼0.5, 1 mM IPTG was added to induce the TAT substrates. The cells were harvested at 2,400 x g for 5 min and washed with 5 mL of 0.75 M sucrose, 10 mM Tris-HCl (pH 7.5). They were then suspended in 1 mL of 0.75 M sucrose, 10 mM Tris-HCl (pH 7.5) and 20 µg/mL lysozyme solution and incubated on ice for 5 min. They were converted into spheroplasts by adding 1.5 mM EDTA (pH 8.0) at equal intervals over 10 min, bringing the total volume to 2 mL. An aliquot of the spheroplast suspension (1 mL) was centrifuged (15,000 x g, 4°C, 1 min) to sediment spheroplasts. The spheroplasts were suspended in 500 µL water and treated with 10% TCA, followed by incubation on ice for 10 min to precipitate proteins. The periplasmic fraction, obtained by sedimentation of the spheroplasts, was treated similarly. The proteins were then recovered by centrifugation (10,000 x g, 4°C, 5 min), washed with acetone, and subjected to SDS-PAGE/immunoblotting, as previously described ^77^.

### Fluorescence microscopy

*E. coli* strains were grown in LB medium to OD_600 nm_ ∼0.5, followed by the addition of 1 mM IPTG to induce TorA-GFP. The cells were fixed on glass slides coated with 1% agarose, as described ^39^. The cells were observed using a fluorescence microscope (Olympus BX53, combined with a U-FGFP filter). Images were captured with a 3CCD digital camera (Olympus DP74). The digital images were then processed and analyzed using the cell image analysis software, cellSens Standard (Olympus).

### Assaying of the protein translocation *in vitro*

The TAT substrates were synthesized *in vitro* using the *E. coli* T7 S30 extract system. For the co-translational translocation assay, the followings were added to a 20 µL reaction mixture: [^35^S] methionine and cysteine (∼10 MBq/mL), plasmids encoding the substrate protein under the T7 promoter, and INV or proteoliposomes. The synthesis/translocation of the RI-labeled substrates were performed at 37°C for 1 h. When INV was used, it was added to the reaction mixture 5 min after the reaction started. For the post-translational translocation assay, the RI-labeled substrates (3 µL), synthesized in the absence of INV or proteoliposomes at 30°C for 30 min, were added to a translocation mixture containing 50 mM HEPES-KOH (pH 7.5), INV or proteoliposomes, 1 mM MgSO_4_, 5 mM creatine phosphate, 10 µg/mL creatine kinase, and 1 mM ATP. After incubating the mixture at 37°C for 45 min, it was divided into two portions. One portion was treated with 5% TCA to precipitate the substrate proteins. The other portion was mixed with an equal volume of PK (1 mg/mL) and incubated at 25°C for 20 min. After adding 5% TCA, the samples were incubated at 56°C for 5 min to inactivate PK. Then, the proteins were recovered by centrifugation (10,000 x g, 4°C, 5 min). Radioactive materials were detected and quantified as described above.

### TALON affinity chromatography

INV (4 mg protein) was solubilized using 1% DDM, 20% glycerol, 50 mM HEPES-KOH (pH 7.5). The solubilized membranes were recovered by centrifugation (170,000 x g, 4°C, 30 min). Samples were applied to a TALON column (0.5 mL). After washing the column with 20 mL of 0.02% DDM, 10% glycerol, and 50 mM HEPES-KOH (pH 7.5), the column-bound materials were eluted with the same buffer containing 150 mM imidazole. The eluted fraction was analyzed by TLC using a solvent system C (chloroform/ethanol/water: 3/7/4) for development. Then, MPIase was detected using anti-MPIase antibody, as described ^53^.

### Reconstitution of proteoliposomes

Phospholipids (13 mg) were resuspended in buffer A (50 mM HEPES-KOH (pH 7.5), 1 mM dithiothreitol) at 20 mg/mL, followed by bath sonication to form liposomes. The liposomes were solubilized in 5% [w/v] OG and incubated on ice for 15 min. Then, 200 mg of Bio Beads SM-2 were added, and the mixture was gently stirred at 4°C for 30 min. Purified TatA (100 µg or 1000 µg), TatB (100 µg), TatC (100 µg) and F_0_F_1_-ATPase (1 µM) were added to the solution. The solution was gently mixed at 4°C for at least 3 h. The Bio Beads SM-2 were then removed, and the proteoliposomes were recovered by centrifugation (170,000 x g, 4°C, 1 h). DAG- and DAG/MPIase-containing liposomes were prepared, as described ^77^. TatABC/F_0_F_1_-ATPase proteoliposomes and PL/DAG liposomes or PL/DAG/MPIase liposomes were typically mixed at a ratio of 4:1, followed by three cycles of freezing, thawing, and sonicating to allow liposome fusion, as described ^54^. As a result of liposome fusion, proteoliposomes contain DAG and MPIase at 5% each to phospholipids. The mixture was then passed through a mini-extruder (Avanti Polar Lipids, Inc.) with a 0.4 µm pore size filter to obtain uniformly sized liposomes, as described ^64^.

### Subcellular fractionation

After the cells were grown in LB medium to OD_600 nm_ ∼0.5, 1 mM IPTG was added to induce the TAT substrates. The harvested cells were suspended in 50 mM HEPES-KOH (pH 7.5), 10% glycerol, 1 mM dithiothreitol, 1 mM phenylmethylsulfonyl fluoride, protease inhibitor cocktail (Roche) and then disrupted using a French press. Then, the cells were incubated with 2 mM MgSO_4_, DNase I (1 mg/mL) on ice for 15 min, followed by the addition of 2 mM EDTA (pH 8.0) and incubation on ice for an additional 15 min. The disrupted and undisrupted cells were separated by centrifugation (15,000 x g, 4°C, 10 min) and then they were fractionated into soluble (cytosol and periplasm) and insoluble (membrane) fractions by centrifugation (150,000 x g, 4°C, 1 h). TorA-GFP was detected by monitoring GFP fluorescence. Samples were added to 2 mL of 50 mM HEPES-KOH (pH 7.5), and the fluorescence intensity was measured at an excitation of 395 nm and an emission of 509 nm using an FP-8200 fluorescence spectrometer (JASCO).

### Statistical analysis

Statistical significance was determined using a Student’s t-test. The *P* values of less than 0.05 were considered significant.

### Other methods

SDS-PAGE was performed using 12.5% acrylamide/0.27% *N*,*N*’-bis (methyleneacrylamide) ^78^ for TatA, TatB, TatC, SecB, β-lactamase, and MPIase. 15.0% acrylamide/0.33% *N*,*N*’-bis (methyleneacrylamide) ^79^ was used to analyze SufI and TorA-GFP, while 8% acrylamide/0.17% *N*,*N*’-bis (methyleneacrylamide) was used for TorA. Bands on the immunoblots were quantified using a CS analyzer (ATTO). Protein quantification after INV preparation was performed using bovine serum albumin as a standard, as described ^80^. Detection of MPIase and other proteins by immunoblotting was performed as described ^58,81^.

## Supporting information

Source data for GFP assay in Fig. 4A

## Acknowledgments

We thank Prof. Matthias Müller for the fruitful discussions and antibodies; Dr. Yoshimi Kikuchi for the discussions and plasmids encoding TorA-GFP; and Ms. Hiromi Saito, Mari Saikudo and Miki Sawaguchi for the MPIase purification. We also thank NBRP (NIG, Japan): *E. coli* for the Keio clones. Experiments involving radioisotopes were carried out in the RI Laboratory at Iwate University.

## Author contributions

Conceptualization: HN, KN

Methodology: HN, KS, MY, KN

Investigation: HN, NY, HK, YS, KK

Visualization: HN, NY, YS, KK, KN

Funding acquisition: HN, KN

Project administration: HN, KN

Supervision: HN, MY, KN

Writing – original draft: HN, KN

Writing – review & editing: HN, NY, YS, KK, KS, MY, KN

## Data availability

Source Data underlying the graphs in the manuscript can be found in Supplementary Data 1. Full, unedited immunoblots and autoradiograms are shown in Fig. S11. All other data are available from the corresponding author on reasonable request.

## References

1. Chaddock, A. M. et al. A new type of signal peptide: central role of a twin-arginine motif in transfer signals for the ΔpH-dependent thylakoidal protein translocase. EMBO J. 14, 2715–2722 (1995).

2. Mould, R. M. & Robinson, C. A proton gradient is required for the transport of two lumenal oxygen-evolving proteins across the thylakoid membrane. J. Biol. Chem. 266, 12189–12193 (1991).

3. Berks, B. C. The twin-arginine protein translocation pathway. Annu. Rev. Biochem. 84, 843–864 (2015).

4. Wu, L. F., Ize, B., Chanal, A., Quentin, Y. & Fichant, G. Bacterial twin-arginine signal peptide-dependent protein translocation pathway: evolution and mechanism. J. Mol. Microbiol. Biotechnol. 2, 179–189 (2000).

5. Berks, B. C. A common export pathway for proteins binding complex redox cofactors? Mol. Microbiol. 22, 393–404 (1996).

6. Cristóbal, S., de Gier, J. W., Nielsen, H. & von Heijne, G. Competition between Sec- and TAT-dependent protein translocation in *Escherichia coli*. EMBO J. 18, 2982–2990 (1999).

7. Blaudeck, N., Kreutzenbeck, P., Freudl, R. & Sprenger, G. A. Genetic analysis of pathway specificity during posttranslational protein translocation across the *Escherichia coli* plasma membrane. J. Bacteriol. 185, 2811–2819 (2003).

8. Bogsch, E., Brink, S. & Robinson, C. Pathway specificity for a ΔpH-dependent precursor thylakoid lumen protein is governed by a ‘Sec-avoidance’ motif in the transfer peptide and a ‘Sec-incompatible’ mature protein. EMBO J. 16, 3851–3859 (1997).

9. Palmer, T., Sargent, F. & Berks, B. C. Export of complex cofactor-containing proteins by the bacterial Tat pathway. Trends Microbiol. 13, 175–180 (2005).

10. Yahr, T. L. & Wickner, W. T. Functional reconstitution of bacterial Tat translocation *in vitro*. EMBO J. 20, 2472–2479 (2001).

11. Bageshwar, U. K. & Musser, S. M. Two electrical potential-dependent steps are required for transport by the *Escherichia coli* Tat machinery. J. Cell Biol. 179, 87–99 (2007).

12. Zhou, W., Hao, B., Bricker, T. M. & Theg, S. M. A real-time analysis of protein transport via the twin arginine translocation pathway in response to different components of the protonmotive force. J. Biol. Chem. 299, 105286 (2023).

13. De Leeuw, E., Porcelli, I., Sargent, F., Palmer, T. & Berks, B. C. Membrane interactions and self-association of the TatA and TatB components of the twin-arginine translocation pathway. FEBS Lett. 506, 143–148 (2001).

14. Sargent, F. et al. Overlapping functions of components of a bacterial Sec-independent protein export pathway. EMBO J. 17, 3640–3650 (1998).

15. Weiner, J. H. et al. A novel and ubiquitous system for membrane targeting and secretion of cofactor-containing proteins. Cell 93, 93–101 (1998).

16. Bogsch, E. G. et al. An essential component of a novel bacterial protein export system with homologues in plastids and mitochondria. J. Biol. Chem. 273, 18003–18006 (1998).

17. Jack, R. L., Sargent, F., Berks, B. C., Sawers, G. & Palmer, T. Constitutive expression of *Escherichia coli* tat genes indicates an important role for the twin-arginine translocase during aerobic and anaerobic growth. J. Bacteriol. 183, 1801–1804 (2001).

18. Sargent, F., Stanley, N. R., Berks, B. C. & Palmer, T. Sec-independent protein translocation in *Escherichia coli*. A distinct and pivotal role for the TatB protein. J. Biol. Chem. 274, 36073–36082 (1999).

19. Hicks, M. G. et al. The *Escherichia coli* twin-arginine translocase: conserved residues of TatA and TatB family components involved in protein transport. FEBS Lett. 539, 61–67 (2003).

20. Yen, M.-R., Tseng, Y.-H., Nguyen, E. H., Wu, L.-F. & Saier, M. H. Sequence and phylogenetic analyses of the twin-arginine targeting (Tat) protein export system. Arch. Microbiol. 177, 441–450 (2002).

21. Walther, T. H., Grage, S. L., Roth, N. & Ulrich, A. S. Membrane alignment of the pore-forming component TatA(d) of the twin-arginine translocase from *Bacillus subtilis* resolved by solid-state NMR spectroscopy. J. Am. Chem. Soc. 132, 15945–15956 (2010).

22. Rodriguez, F., et al. Structural model for the protein-translocating element of the twin-arginine transport system. Proc. Natl. Acad. Sci. U S A 110, E1092–E1101 (2013).

23. Rollauer, S. E. et al. Structure of the TatC core of the twin-arginine protein transport system. Nature 492, 210–214 (2012).

24. Ramasamy, S., Abrol, R., Suloway, C. J. M. & Clemons, W. M. The glove-like structure of the conserved membrane protein TatC provides insight into signal sequence recognition in twin-arginine translocation. Structure 21, 777–788 (2013).

25. Bolhuis, A., Mathers, J. E., Thomas, J. D., Barrett, C. M. & Robinson, C. TatB and TatC form a functional and structural unit of the twin-arginine translocase from *Escherichia coli*. J. Biol. Chem. 276, 20213–20219 (2001).

26. Cline, K. & Mori, H. Thylakoid ΔpH-dependent precursor proteins bind to a cpTatC-Hcf106 complex before Tha4-dependent transport. J. Cell Biol. 154, 719–729 (2001).

27. de Leeuw, E. et al. Oligomeric properties and signal peptide binding by *Escherichia coli* Tat protein transport complexes. J. Mol. Biol. 322, 1135–1146 (2002).

28. Alami, M. et al. Differential interactions between a twin-arginine signal peptide and its translocase in *Escherichia coli*. Mol. Cell 12, 937–946 (2003).

29. Richter, S. & Brüser, T. Targeting of unfolded PhoA to the TAT translocon of *Escherichia coli*. J. Biol. Chem. 280, 42723–42730 (2005).

30. Tarry, M. J. et al. Structural analysis of substrate binding by the TatBC component of the twin-arginine protein transport system. Proc. Natl. Acad. Sci. U S A 106, 13284–13289 (2009).

31. Dabney-Smith, C. & Cline, K. Clustering of C-terminal stromal domains of Tha4 homo-oligomers during translocation by the Tat protein transport system. Mol. Biol. Cell 20, 2060–2069 (2009).

32. Leake, M. C. et al. Variable stoichiometry of the TatA component of the twin-arginine protein transport system observed by *in vivo* single-molecule imaging. Proc. Natl. Acad. Sci. U S A 105, 15376–15381 (2008).

33. Gohlke, U. et al. The TatA component of the twin-arginine protein transport system forms channel complexes of variable diameter. Proc. Natl. Acad. Sci. U S A 102, 10482–10486 (2005).

34. Beck, D. et al. Ultrastructural characterisation of *Bacillus subtilis* TatA complexes suggests they are too small to form homooligomeric translocation pores. Biochim. Biophys. Acta 1833, 1811–1819 (2013).

35. McDevitt, C. A., Buchanan, G., Sargent, F., Palmer, T. & Berks, B. C. Subunit composition and *in vivo* substrate-binding characteristics of *Escherichia coli* Tat protein complexes expressed at native levels. FEBS J. 273, 5656–5668 (2006).

36. Oates, J. et al. The *Escherichia coli* twin-arginine translocation apparatus incorporates a distinct form of TatABC complex, spectrum of modular TatA complexes and minor TatAB complex. J. Mol. Biol. 346, 295–305 (2005).

37. Mori, H. & Cline, K. A twin arginine signal peptide and the pH gradient trigger reversible assembly of the thylakoid ΔpH/Tat translocase. J. Cell Biol. 157, 205–210 (2002).

38. Alcock, F., et al. Live cell imaging shows reversible assembly of the TatA component of the twin-arginine protein transport system. Proc. Natl. Acad. Sci. U S A 110, E3650–E3659 (2013).

39. Rose, P., Fröbel, J., Graumann, P. L. & Müller, M. Substrate-dependent assembly of the Tat translocase as observed in live *Escherichia coli* cells. PLoS One 8, e69488 (2013).

40. Dabney-Smith, C., Mori, H. & Cline, K. Oligomers of Tha4 organize at the thylakoid Tat translocase during protein transport. J. Biol. Chem. 281, 5476–5483 (2006).

41. Brüser, T. & Sanders, C. An alternative model of the twin arginine translocation system. Microbiol. Res. 158, 7–17 (2003).

42. Mehner-Breitfeld, D. et al. TatA and TatB generate a hydrophobic mismatch important for the function and assembly of the Tat translocon in *Escherichia coli*. J. Biol. Chem. 298, 102236 (2022).

43. Lüke, I., Handford, J. I., Palmer, T. & Sargent, F. Proteolytic processing of *Escherichia coli* twin-arginine signal peptides by LepB. Arch. Microbiol. 191, 919–925 (2009).

44. Cline, K. & McCaffery, M. Evidence for a dynamic and transient pathway through the TAT protein transport machinery. EMBO J. 26, 3039–3049 (2007).

45. Settles, A. M. et al. Sec-independent protein translocation by the maize Hcf106 protein. Science 278, 1467–1470 (1997).

46. Nishiyama, K. et al. MPIase is a glycolipozyme essential for membrane protein integration. Nat. Commun. 3, 1260 (2012).

47. Mori, S. et al. Intermolecular interactions between a membrane protein and a glycolipid essential for membrane protein integration. ACS Chem. Biol. 17, 609–618 (2022).

48. Fujikawa, K. et al. Structural requirements of a glycolipid MPIase for membrane protein integration. Chem. Eur. J. 29, e202300437 (2023).

49. Nomura, K. et al. Alteration of membrane physicochemical properties by two factors for membrane protein integration. Biophys. J. 117, 99–110 (2019).

50. Nishiyama, K. et al. A novel complete reconstitution system for membrane integration of the simplest membrane protein. Biochem. Biophys. Res. Commun. 394, 733–736 (2010).

51. Nishikawa, H., Sasaki, M. & Nishiyama, K. Membrane insertion of F_0_ c subunit of F_0_F_1_ ATPase depends on glycolipozyme MPIase and is stimulated by YidC. Biochem. Biophys. Res. Commun. 487, 477–482 (2017).

52. Moser, M., Nagamori, S., Huber, M., Tokuda, H. & Nishiyama, K. Glycolipozyme MPIase is essential for topology inversion of SecG during preprotein translocation. Proc. Natl. Acad. Sci. U S A 110, 9734–9739 (2013).

53. Endo, Y., Shimizu, Y., Nishikawa, H., Sawasato, K. & Nishiyama, K. Interplay between MPIase, YidC, and PMF during Sec-independent insertion of membrane proteins. Life Sci. Alliance 5, e202101162 (2021).

54. Nishiyama, K., Suzuki, T. & Tokuda, H. Inversion of the membrane topology of SecG coupled with SecA-dependent preprotein translocation. Cell 85, 71–81 (1996).

55. Nishiyama, K. et al. A derivative of lipid A is involved in signal recognition particle/SecYEG-dependent and -independent membrane integrations. J. Biol. Chem. 281, 35667–35676 (2006).

56. Nishikawa, H. et al. Interaction between glycolipid MPIase and proteinaceous factors during protein integration into the cytoplasmic membrane of *E. coli*. Front. Mol. Biosci. 9, 986602 (2022).

57. Sasaki, M. et al. The bacterial protein YidC accelerates MPIase-dependent integration of membrane proteins. J. Biol. Chem. 294, 18898–18908 (2019).

58. Sawasato, K. et al. CdsA is involved in biosynthesis of glycolipid MPIase essential for membrane protein integration *in vivo*. Sci. Rep. 9, 1372 (2019).

59. Alami, M., Trescher, D., Wu, L.-F. & Müller, M. Separate analysis of twin-arginine translocation (Tat)-specific membrane binding and translocation in *Escherichia coli*. J. Biol. Chem. 277, 20499– 20503 (2002).

60. Santini, C. L. et al. A novel sec-independent periplasmic protein translocation pathway in *Escherichia coli*. EMBO J. 17, 101–112 (1998).

61. Stiegler, N., Dalbey, R. E. & Kuhn, A. M13 procoat protein insertion into YidC and SecYEG proteoliposomes and liposomes. J. Mol. Biol. 406, 362–370 (2011).

62. Majdalani, N. & Ippen-Ihler, K. Membrane insertion of the F-pilin subunit is Sec independent but requires leader peptidase B and the proton motive force. J. Bacteriol. 178, 3742–3747 (1996).

63. Rathmann, C., Schlösser, A. S., Schiller, J., Bogdanov, M. & Brüser, T. Tat transport in *Escherichia coli* requires zwitterionic phosphatidylethanolamine but no specific negatively charged phospholipid. FEBS Lett. 591, 2848–2858 (2017).

64. Kawashima, Y., Miyazaki, E., Müller, M., Tokuda, H. & Nishiyama, K. Diacylglycerol specifically blocks spontaneous integration of membrane proteins and allows detection of a factor-assisted integration. J. Biol. Chem. 283, 24489–24496 (2008).

65. Shanmugham, A., Wong Fong Sang, H. W., Bollen, Y. J. M. & Lill, H. Membrane binding of twin arginine preproteins as an early step in translocation. Biochemistry 45, 2243–2249 (2006).

66. Mikhaleva, N. I., Santini, C. L., Giordano, G., Nesmeyanova, M. A. & Wu, L. F. Requirement for phospholipids of the translocation of the trimethylamine N-oxide reductase through the Tat pathway in *Escherichia coli*. FEBS Lett. 463, 331–335 (1999).

67. Sikdar, R. & Doerrler, W. T. Inefficient Tat-dependent export of periplasmic amidases in an *Escherichia coli* strain with mutations in two DedA family genes. J. Bacteriol. 192, 807–818 (2010).

68. Bogdanov, M. & Dowhan, W. Phospholipid-assisted protein folding: phosphatidylethanolamine is required at a late step of the conformational maturation of the polytopic membrane protein lactose permease. EMBO J. 17, 5255–5264 (1998).

69. Ma, X. & Browse, J. Altered rates of protein transport in *Arabidopsis* mutants deficient in chloroplast membrane unsaturation. Phytochemistry 67, 1629–1636 (2006).

70. Sawasato, K., Sekiya, Y. & Nishiyama, K. Two-step induction of *cdsA* promoters leads to upregulation of the glycolipid MPIase at cold temperature. FEBS Lett. 593, 1711–1723 (2019).

71. Hikage, R., Tadika, Y., Asanuma, H., Han, Y. & Nishiyama, K. MucA is a small peptide encoded by an overlapping sequence with *cdsA* that upregulates the biosynthesis of glycolipid MPIase in the cold. Biochem. Biophys. Res. Commun. 721, 150148 (2024).

72. Kikuchi, Y., Date, M., Itaya, H., Matsui, K. & Wu, L.-F. Functional analysis of the twin-arginine translocation pathway in *Corynebacterium glutamicum* ATCC 13869. Appl. Environ. Microbiol. 72, 7183–7192 (2006).

73. Kikuchi, Y., Itaya, H., Date, M., Matsui, K. & Wu, L.-F. TatABC overexpression improves *Corynebacterium glutamicum* Tat-dependent protein secretion. *App*. Environ. Microbiol. 75, 603– 607 (2009).

74. Umakoshi, M. et al. Improving protein secretion of a transglutaminase-secreting *Corynebacterium glutamicum* recombinant strain on the basis of ^13^C metabolic flux analysis. J. Biosci. Bioeng. 112, 595–601 (2011).

75. Cline, K., Ettinger, W. F. & Theg, S. M. Protein-specific energy requirements for protein transport across or into thylakoid membranes. Two lumenal proteins are transported in the absence of ATP. J. Biol. Chem. 267, 2688–2696 (1992).

76. Suzuki, T., Ueno, H., Mitome, N., Suzuki, J. & Yoshida, M. F_0_ of ATP synthase is a rotary proton channel. Obligatory coupling of proton translocation with rotation of c-subunit ring. J. Biol. Chem. 277, 13281–13285 (2002).

77. Nishikawa, H., Sasaki, M. & Nishiyama, K. *In vitro* assay for bacterial membrane protein integration into proteoliposomes. Bio Protoc. 10, e3626 (2020).

78. Hussain, M., Ichihara, S. & Mizushima, S. Accumulation of glyceride-containing precursor of the outer membrane lipoprotein in the cytoplasmic membrane of *Escherichia coli* treated with globomycin. J. Biol. Chem. 255, 3707–3712 (1980).

79. Laemmli, U. K. Cleavage of structural proteins during the assembly of the head of bacteriophage T4. Nature 227, 680–685 (1970).

80. Lowry, O. H., Rosebrough, N. J., Farr, A. L. & Randall, R. J. Protein measurement with the Folin phenol reagent. J. Biol. Chem. 193, 265–275 (1951).

81. Ikegami, A., Nishiyama, K., Matsuyama, S. & Tokuda, H. Disruption of *rpmJ* encoding ribosomal protein L36 decreases the expression of *secY* upstream of the *spc* operon and inhibits protein translocation in *Escherichia coli*. Biosci. Biotechnol. Biochem. 69, 1595–1602 (2005).

