## Supplementary_information for "Glycolipid MPIase is essential for the TAT (Twin-Arginine Translocation) pathway"

**The PDF file includes:**

Figs. S1 to S11
Tables S1 to S3
Supplementary References (1~8)

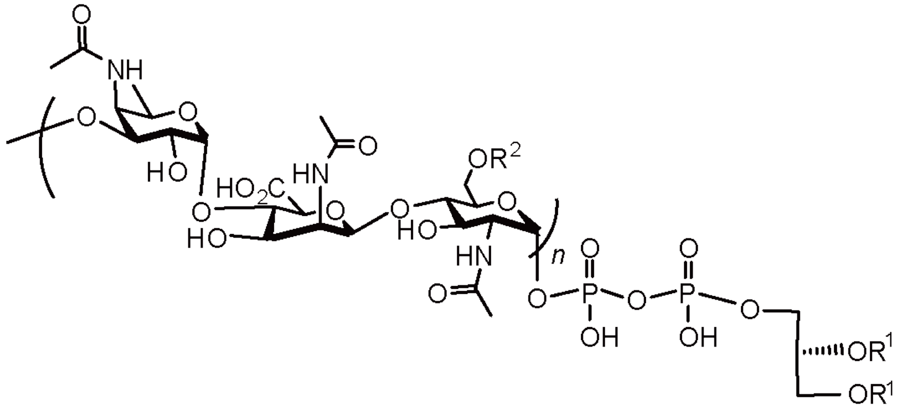

**Fig. S1. Structure of MPIase.** The repeating unit of the glycan chain consists of a trisaccharide unit of 4-acetamido-4-deoxyfucose (Fuc4NAc), 2-acetamido-2-deoxymannuronic acid (ManNAcA), and *N*-acetylglucosamine (GlcNAc). $R^1$: acyl chain of $C_{16}$ or $C_{18}$, $R^2$: Ac or H. n=9~11.

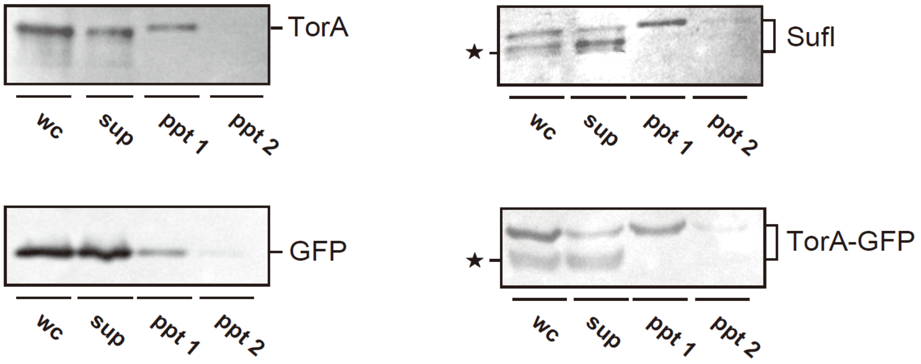


**Fig. S2. The level of inclusion bodies of TAT substrates.** EK413 cells that expressed the specified proteins were disrupted using a French press (whole cells; wc), followed by centrifugation at 15,000 x g for 10 min at 4°C, yielding a pellet (ppt 1). The resulting supernatant was centrifuged again under the same conditions to obtain the supernatant (sup) and a second pellet (ppt2). Samples equivalent to 10 µg of whole-cell proteins were analyzed by SDS-PAGE/immunoblotting. The positions of the specified proteins, including the precursor and mature forms, are shown. Stars denote degradation products.

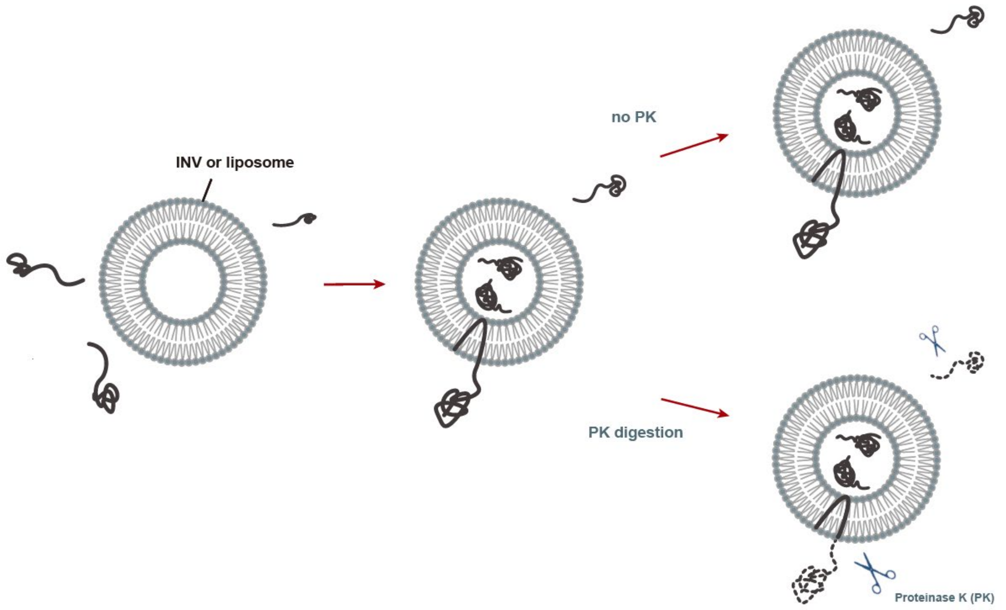


**Fig. S3. Schematic diagram of the *in vitro* protein translocation assay system using INV or proteoliposomes.** Proteins are synthesized *in vitro* in the presence of INV or proteoliposomes. After the translocation reaction, the portion of the protein that has been translocated into the membrane vesicles is protected from cleavage by externally added proteinase K (PK).

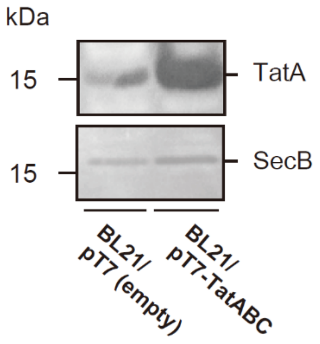


**Fig. S4. Expression levels of TatA in the INV used in Fig. 2A.** Expression levels of TatA in the BL21(DE3)/pT7-TatABC and BL21(DE3)/pT7 (empty) cells (10 µg of protein each) was detected by SDS-PAGE and immunoblotting with an anti-TatA antibody. As a loading control, SecB was detected in the same cells using an anti-SecB antibody.

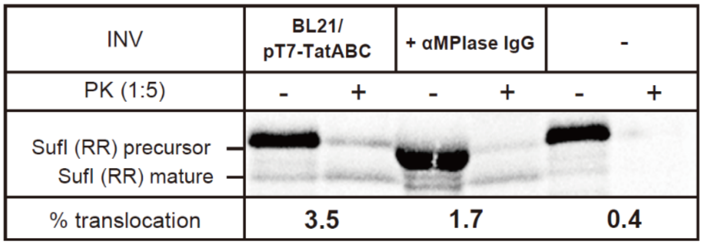


**Fig. S5. The anti-MPIase antibody inhibits SufI translocation into TatABC-overexpressed INV.** INV prepared from BL21/pT7-TatABC (4 µg) and anti-MPIase IgG (αMPIase IgG) (4 µg/µL) were mixed and incubated on ice for 30 min. The translocation assay was performed, as described in the legend to Fig. 2A. Samples were analyzed by SDS-PAGE/autoradiography. One-fifth of the reaction mixture was used as the translation control (-PK). Translocation activity is shown at the bottom of each gel.

.

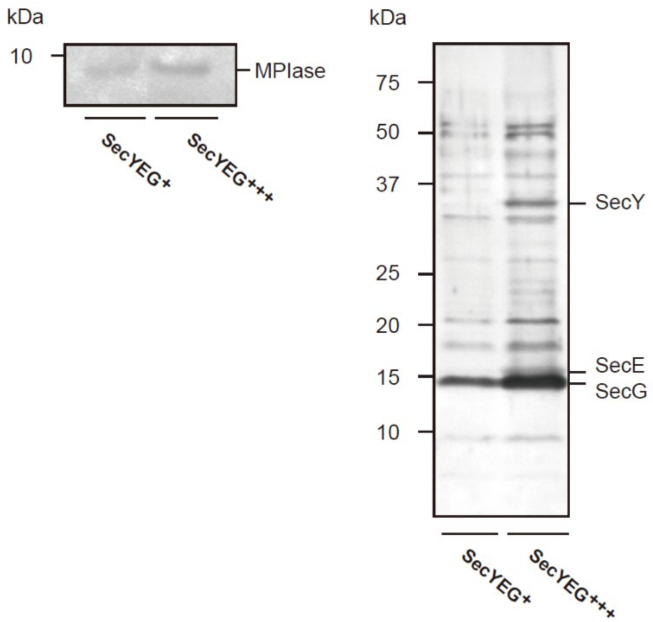


**Fig. S6. SecYEG overproduction does not induce MPIase upregulation.** BL21/pAra-SecYEG was cultivated with or without 0.2% arabinose and then incubated for an additional 1 h. An aliquot (500 µL) of each sample was treated with TCA to a final concentration of 5%. The TCA precipitate was recovered by centrifugation (10,000 × g, 4°C, 5 min) followed by washing with acetone. The samples were then analyzed by SDS-PAGE/immunoblotting using an anti-MPIase antibody (left) or Coomassie Brilliant Blue staining (right).

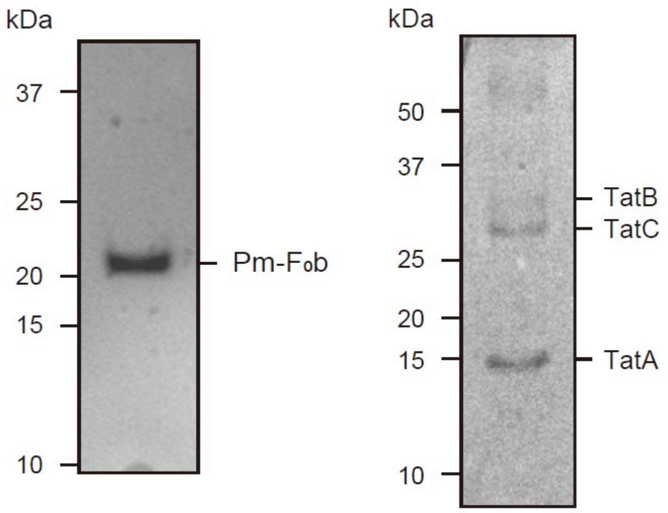


**Fig. S7. Purification of Pm-F$_0$b and TatABC.** BL21(DE3)/pT7-Pm-F$_0$b-His (left) and BL21(DE3)/pT7-TatABC-His (right) were grown in LB medium to OD$_{600nm}$ ~0.5 at 37°C, followed by induction with IPTG (1 mM). INV, then prepared, was solubilized, and then subjected to TALON column chromatography. The eluted fraction (~5 μg) was analyzed by SDS-PAGE/Coomassie Brilliant Blue staining. The positions of the specified proteins are indicated.

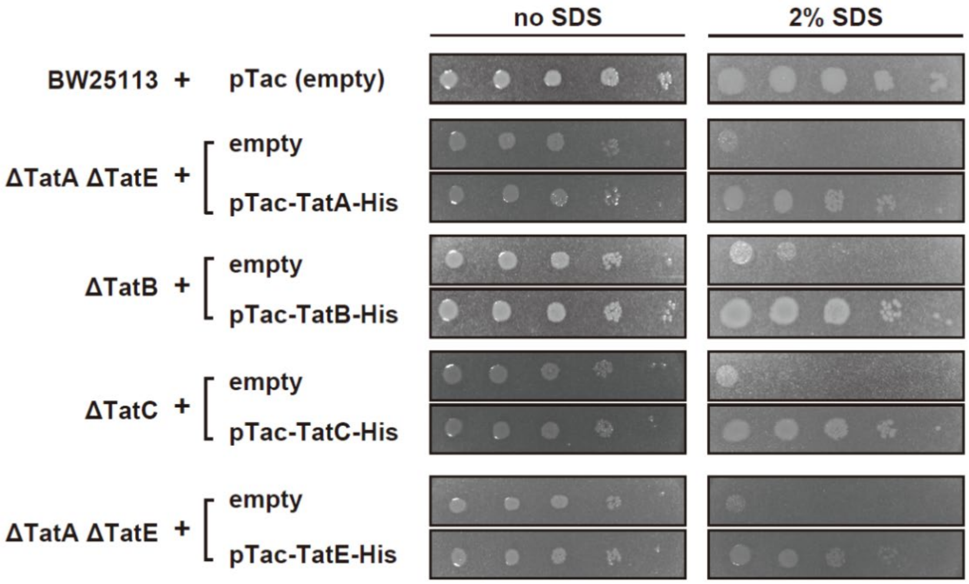


**Fig. S8.** ***In vivo*** **complementation of the SDS-sensitive growth by His-tagged Tat components.** The ΔTatE mutation was transferred to the ΔTatA strain via P1 transduction. Overnight cultures of the ΔTatA/ΔTatE, ΔTatB, and ΔTatC strains harboring the indicated plasmid were adjusted to an $OD_{600nm}$ of ~0.5, serially diluted by a factor of 10, and then spotted on LB agar plates supplemented with or without 2% SDS. All plates were incubated at 37°C for 1 day.

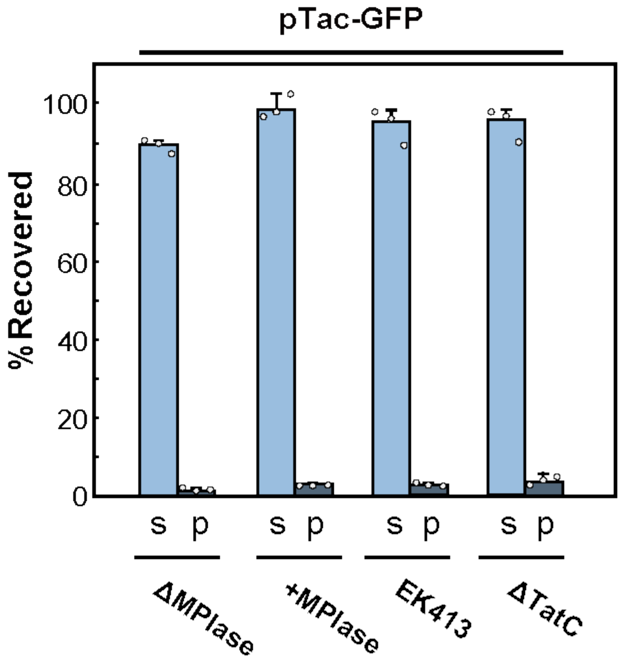


**Fig. S9. GFP without the TAT signal is not targeted to the membrane.** KS23/pTac-GFP cells were cultivated in the presence (+MPIase) or absence (ΔMPIase) of arabinose. EK413/pTac-GFP and JW3815 (ΔTatC)/pTac-GFP were cultivated as well. After inducing GFP with 1 mM IPTG, the cells were disrupted and fractionated to obtain the supernatant (sup; cytosol) and the pellet (ppt; membranes) fractions. The fluorescence intensity of each fraction was measured. The ratio to the whole cell was determined and shown as percentages. The average of three independent experiments, along with the standard deviation, is shown with each value.

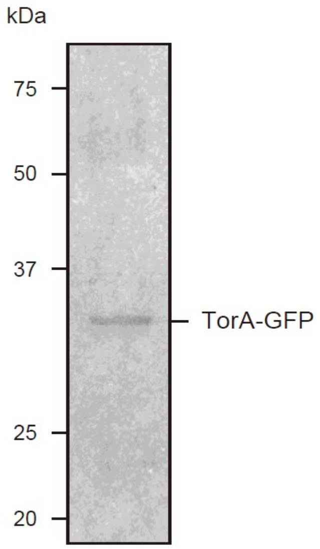


**Fig. S10. Purification of TorA-GFP.** BL21(DE3)/pT7-TorA-GFP was grown in LB medium to $OD_{600nm}$ ~0.5. Then, 1 mM IPTG was added to induce TorA-GFP, and INV was prepared. The solubilized membrane was prepared and then applied to a TALON column. An eluted fraction (5 µL) was subjected to SDS-PAGE and stained with Coomassie Brilliant Blue.

Fig.1A

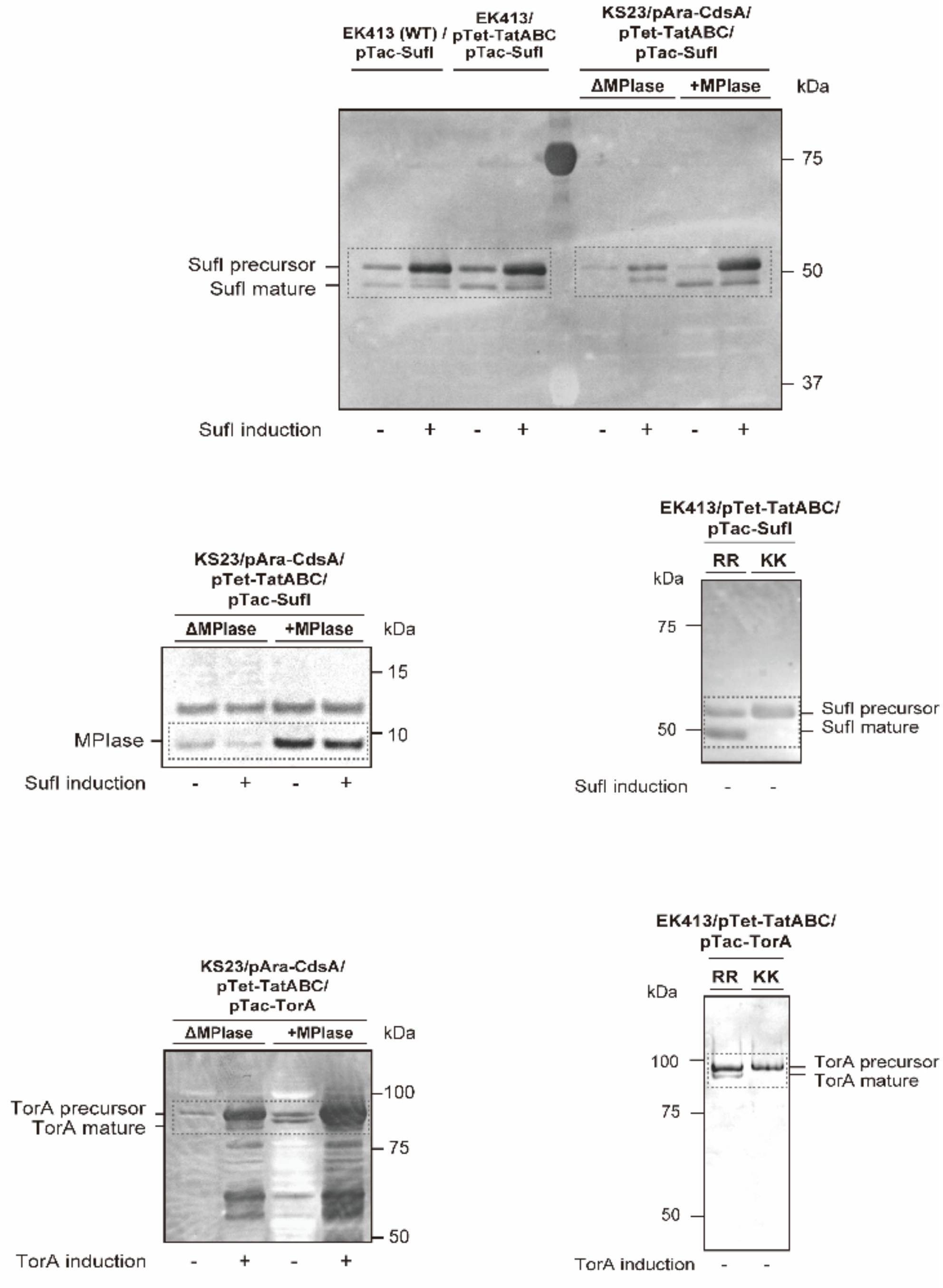

EK413 (WT) / pTac-SufI
EK413/ pTet-TatABC pTac-SufI
KS23/pAra-CdsA/ pTet-TatABC/ pTac-SufI
ΔMPlase
+MPlase
kDa
75
50
37
SufI precursor
SufI mature
SufI induction
- + - + - + - +
KS23/pAra-CdsA/ pTet-TatABC/ pTac-SufI
ΔMPlase
+MPlase
kDa
15
10
MPlase
SufI induction
- + - +
EK413/pTet-TatABC/ pTac-SufI
RR
KK
kDa
75
50
SufI precursor
SufI mature
SufI induction
- -
KS23/pAra-CdsA/ pTet-TatABC/ pTac-TorA
ΔMPlase
+MPlase
kDa
100
75
50
TorA precursor
TorA mature
TorA induction
- + - +
EK413/pTet-TatABC/ pTac-TorA
RR
KK
kDa
100
75
50
TorA precursor
TorA mature
TorA induction
- -

Fig.1B

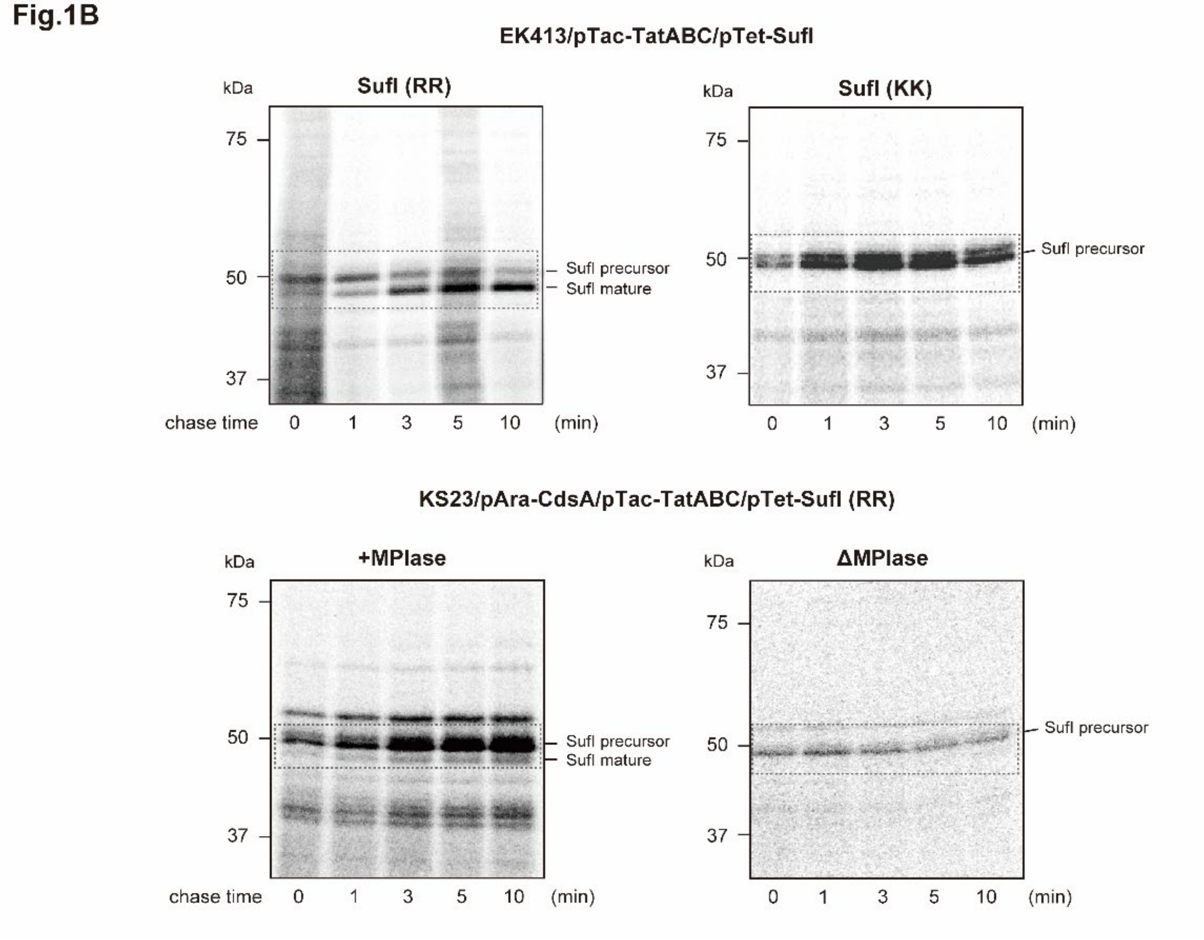
EK413/pTac-TatABC/pTet-SufI
SufI (RR)
SufI (KK)
kDa
75
50
37
SufI precursor
SufI mature
chase time
0
1
3
5
10
(min)
KS23/pAra-CdsA/pTac-TatABC/pTet-SufI (RR)
+MPIase
ΔMPIase

Fig.1C

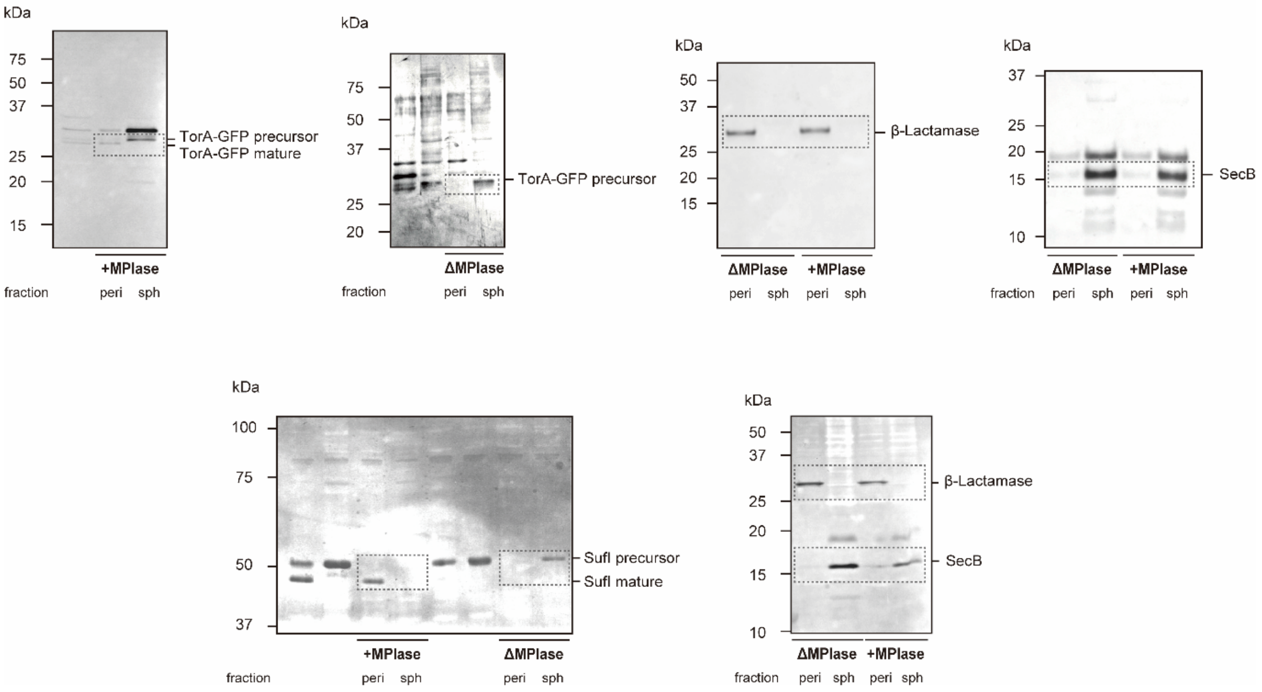
kDa
75
50
37
25
20
15
TorA-GFP precursor
TorA-GFP mature
+MPlase
fraction
peri
sph
kDa
75
50
37
25
20
TorA-GFP precursor
ΔMPlase
fraction
peri
sph
kDa
50
37
25
20
15
β-Lactamase
ΔMPlase
+MPlase
peri
sph
peri
sph
kDa
37
25
20
15
10
SecB
ΔMPlase
+MPlase
fraction
peri
sph
peri
sph
kDa
100
75
50
37
SufI precursor
SufI mature
+MPlase
ΔMPlase
fraction
peri
sph
peri
sph
kDa
50
37
25
20
15
10
β-Lactamase
SecB
ΔMPlase
+MPlase
fraction
peri
sph
peri
sph

Fig.2A

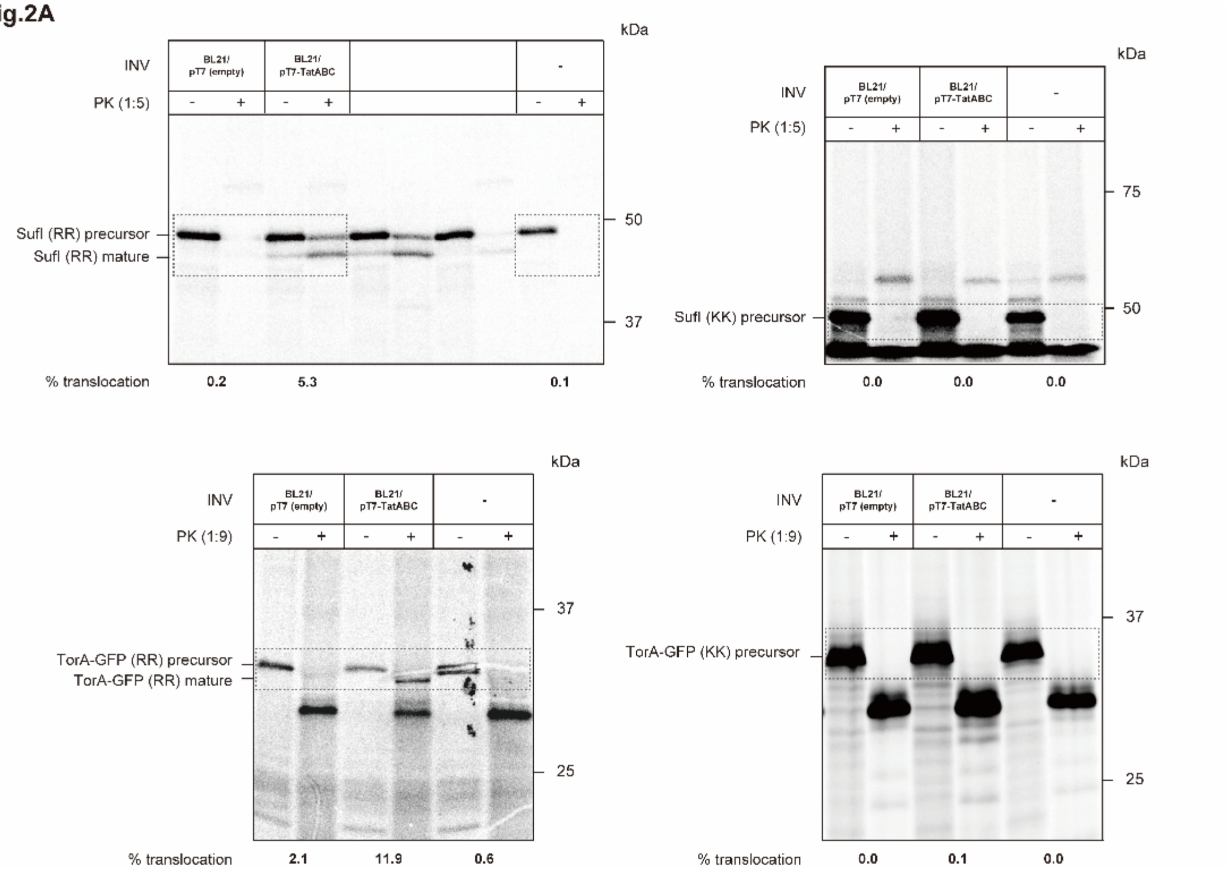


Fig.2B

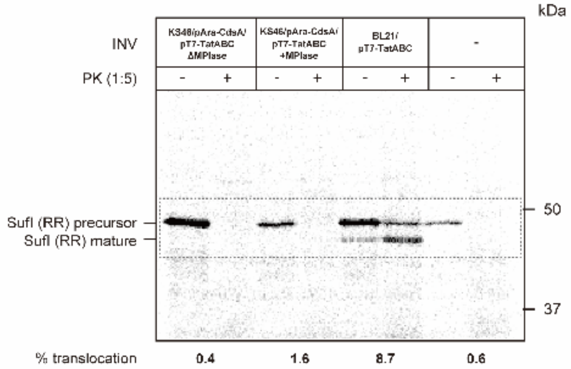


Fig.2C

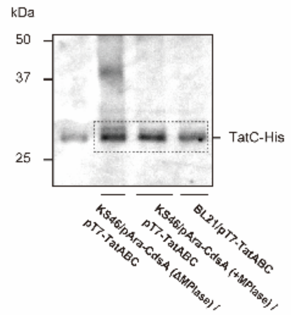


Fig.2D

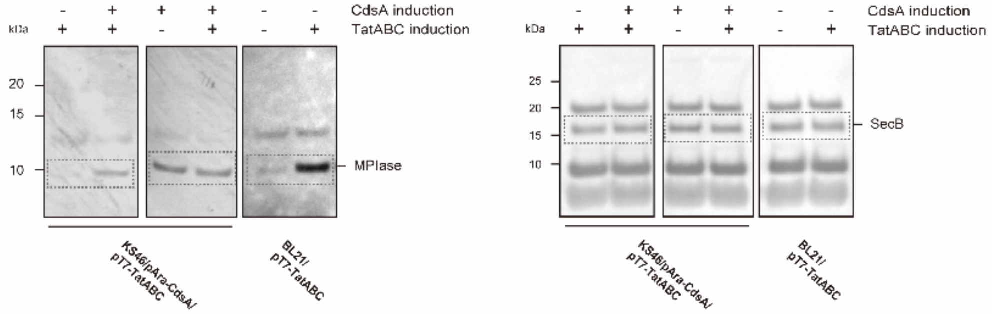

Fig.3A

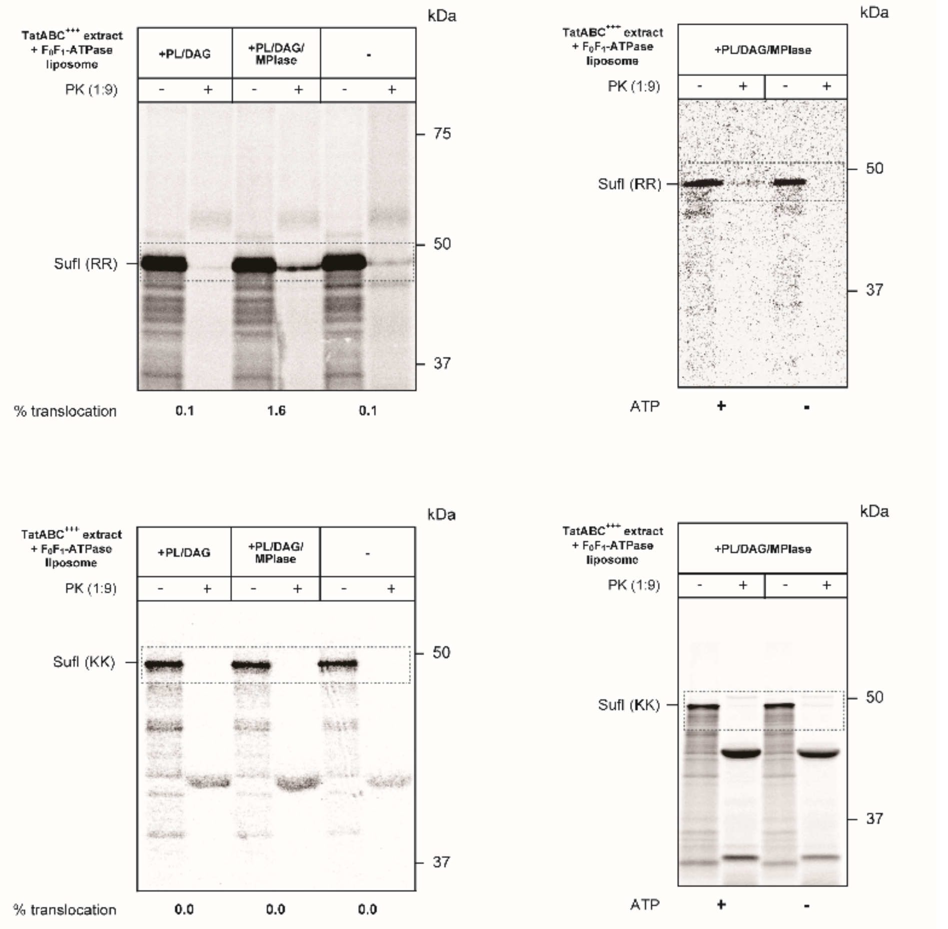
kDa
TatABC+++ extract
+ F0F1-ATPase
liposome
+PL/DAG
+PL/DAG/
MPIase
-
PK (1:9)
- + - + - +
75
50
37
SufI (RR)
% translocation
0.1
1.6
0.1
kDa
TatABC+++ extract
+ F0F1-ATPase
liposome
+PL/DAG/MPIase
PK (1:9)
- + - +
50
37
SufI (RR)
ATP
+
-
kDa
TatABC+++ extract
+ F0F1-ATPase
liposome
+PL/DAG
+PL/DAG/
MPIase
-
PK (1:9)
- + - + - +
50
37
SufI (KK)
% translocation
0.0
0.0
0.0
kDa
TatABC+++ extract
+ F0F1-ATPase
liposome
+PL/DAG/MPIase
PK (1:9)
- + - +
50
37
SufI (KK)
ATP
+
-

Fig.3C

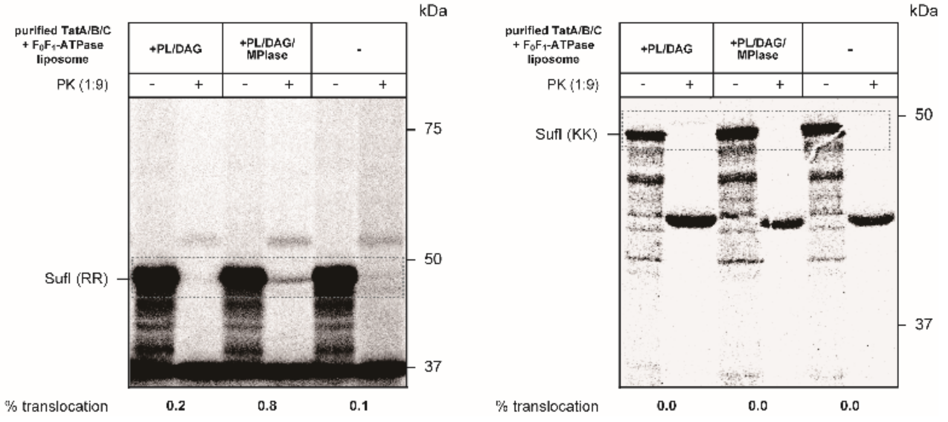
kDa
purified TatA/B/C
+ F0F1-ATPase
liposome
+PL/DAG
+PL/DAG/
MPIase
-
PK (1:9)
- + - + - +
75
50
37
SufI (RR)
% translocation
0.2
0.8
0.1
kDa
purified TatA/B/C
+ F0F1-ATPase
liposome
+PL/DAG
+PL/DAG/
MPIase
-
PK (1:9)
- + - + - +
50
37
SufI (KK)
% translocation
0.0
0.0
0.0

Fig.3D

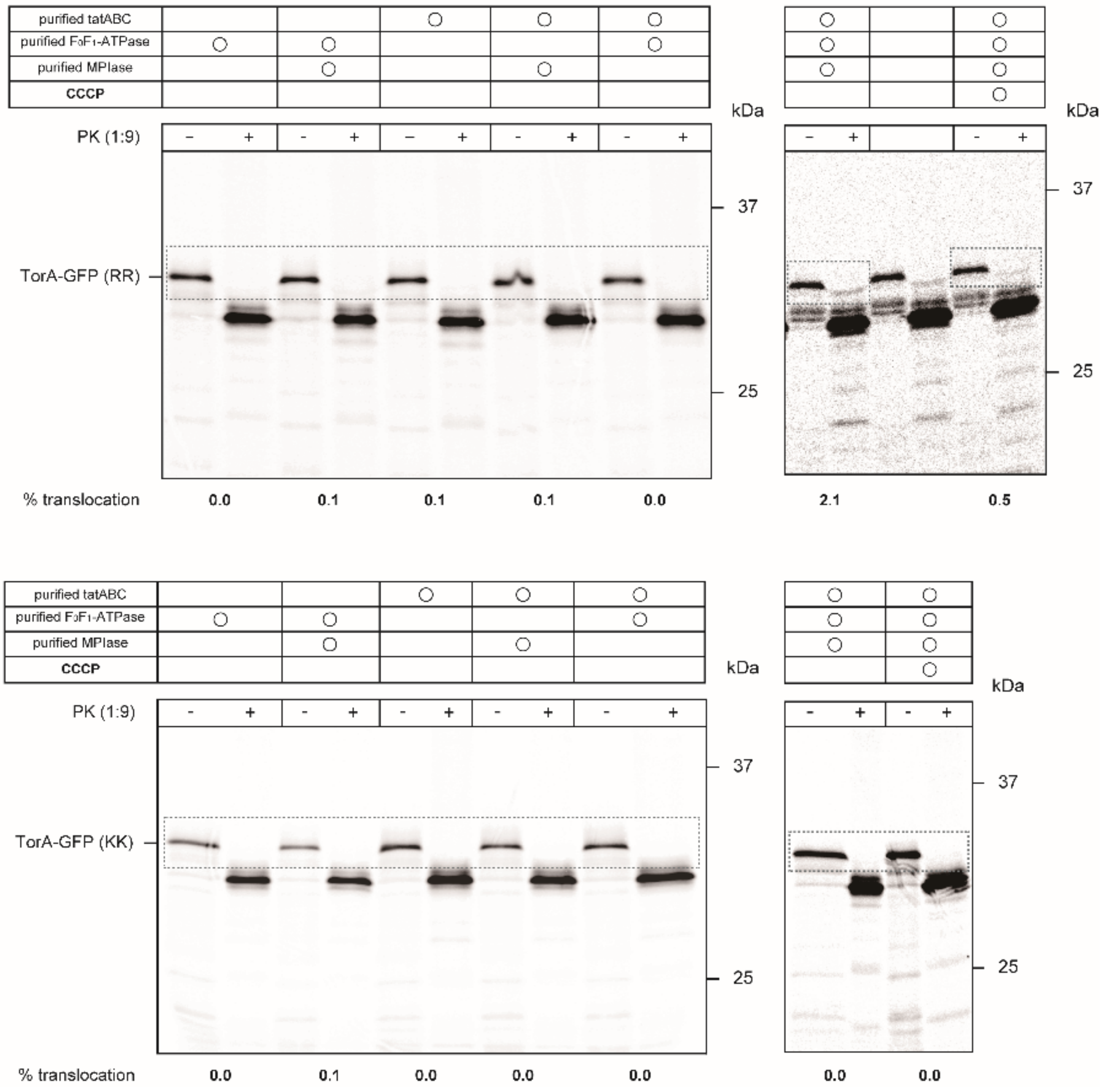

purified tatABC
purified F₀F₁-ATPase
purified MPIase
CCCP
PK (1:9)
kDa
37
25
TorA-GFP (RR)
% translocation
0.0
0.1
0.1
0.1
0.0
2.1
0.5
TorA-GFP (KK)
0.0
0.1
0.0
0.0
0.0
0.0
0.0

**Fig.S2**

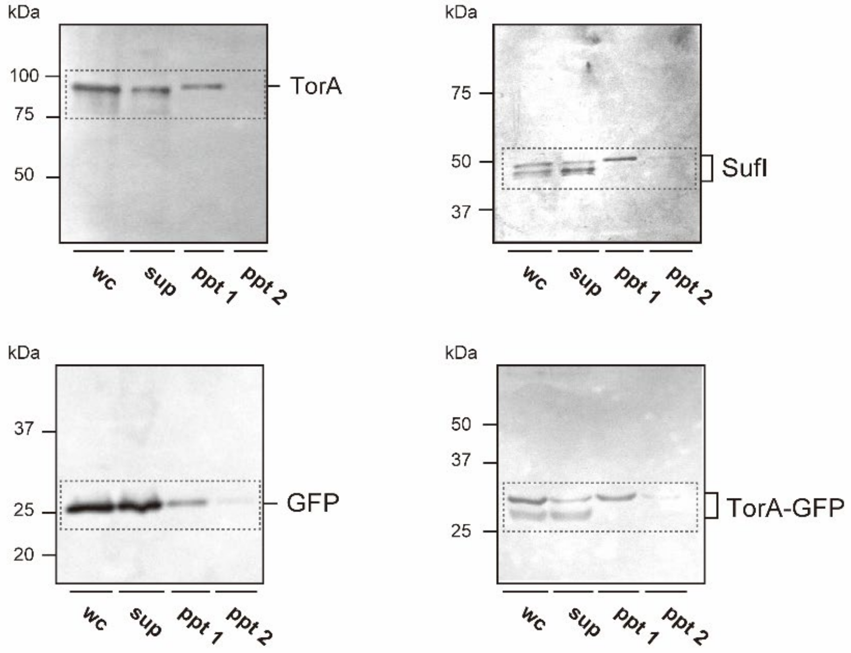

kDa
100
75
50
TorA
wc
sup
ppt 1
ppt 2
kDa
75
50
37
SufI
wc
sup
ppt 1
ppt 2
kDa
37
25
20
GFP
wc
sup
ppt 1
ppt 2
kDa
50
37
25
TorA-GFP
wc
sup
ppt 1
ppt 2

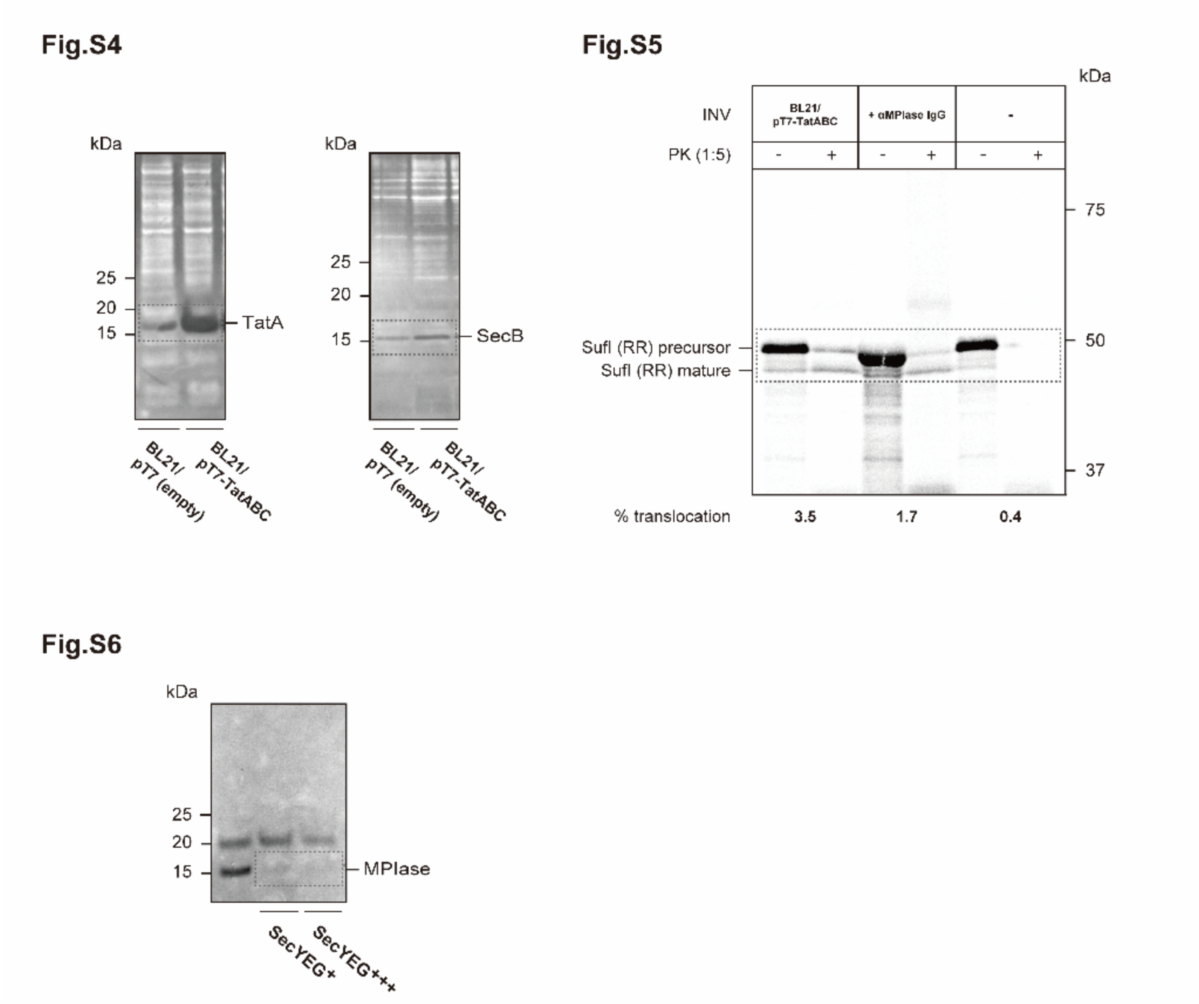


**Fig. S11. Original images used in the figures.**

**Table S1. Strains used in this study.**

| Strains | Relevant genotype and description | Reference |
|---|---|---|
| EK413 | MC4100 *ara*$^{+}$ | [1] |
| DH5α | $F^{-}$, Φ80d*lacZ*ΔM15, Δ(*lacZYA*-*argF*)U169, *deoR*, *recA1*, *endA1*, *hsdR17*($r_k^{-}$, $m_k^{+}$), *phoA*, *supE44*, $\lambda^{-}$, *thi*-1, *gyrA*96, *relA1* | [2] |
| BL21 (DE3) | $F^{-}$, *lon-11*, Δ(*ompT*-*nfrA*)885, Δ(*galM*-*ybhJ*)884, λDE3 [*lacI lacUV5-T7 gene 1 ind1 sam7 nin5*] Δ46 [*mal*$^{+}$]K-12(λS) *hsdS10* | [3] |
| KS23 | EK413 Δ*cdsA*::*cat*, Δ*ynbB* | [4] |
| KS46 | BL21 (DE3) Δ*cdsA*::*cat*, Δ*ynbB*::*kan* | [4] |
| SN1187 | MG1655 Δ*hsdR* Δ*endA* Δ*recA* | [5] |
| BW25113 | $F^{-}$, Δ(*araD-araB*)567 Δ(*rhaD-rhaB*)568 *ΔlacZ*4787 (::rrnB-3) *hsdR514 rph-1* | [6] |
| JW3813 | BW25113 Δ*tatA*::*kan* | [7] |
| JW5580 | BW25113 Δ*tatB*::*kan* | [7] |
| JW3815 | BW25113 Δ*tatC*::*kan* | [7] |
| JW0622 | BW25113 Δ*tatE*::*kan* | [7] |
| HN14 | BW25113 Δ*tatA::kan*, Δ*tatE* | This study |

**Table S2. Plasmids used in this study.**

| Plasmids | Relevant description | Reference |
|---|---|---|
| pUSI2 | Cloning vector containing *tac* promoter, ampicillin resistant | [8] |
| pIVEX-2.3MCS | Cloning vector containing the *T7* promoter, ampicillin resistant | Roche |
| pACYC184-Km | The *cat* gene on pACYC184 was replaced with the *kan* gene. | [4] |
| pETUA | Cloning vector containing the *T7* promoter, ampicillin resistant | BioDynamics Laboratory |
| pTac-SufI-His | $his_6$-*sufI* was cloned into pUSI2 under the control of *tac* promoter | This study |
| pTac-SufI(KK)-His | $his_6$-*sufI(kk)* was cloned into pUSI2 under the control of *tac* promoter | This study |
| pTac-TorA-His | $his_6$-*torA* was cloned into pUSI2 under the control of *tac* promoter | This study |
| pTac-TorA(KK)-His | $his_6$-*torA(kk)* was cloned into pUSI2 under the control of *tac* promoter | This study |
| pTac-TorA-GFP-His | $his_6$-*gfp* with TorA signal sequence was cloned into pUSI2 under the control of *tac* promoter | This study |
| pTac-TorA-GFP(KK)-His | $his_6$-*gfp* with TorA(KK) signal sequence was cloned into pUSI2 under the control of *tac* promoter | This study |
| pTac-TatABC | *tatABC* was cloned into pUSI2 under the control of *tac* promoter | This study |
| pTet-TatABC | *tatABC* was cloned into pACYC-184-Km under the control of *tet* promoter | This study |
| pTet-SufI-His | $his_6$-*sufI* was cloned into pACYC184-Km under the control of *tet* promoter | This study |
| pTac-TatA-His | $his_6$-*tatA* was cloned into pUSI2 under the control of *tac* promoter | This study |
| pTac-TatB-His | $his_6$-*tatB* was cloned into pUSI2 under the control of *tac* promoter | This study |
| pTac-TatC-His | $his_6$-*tatC* was cloned into pUSI2 under the control of *tac* promoter | This study |
| pT7-TatABC | *tatABC* was cloned into pIVEX-2.3MCS under the control of *T7* promoter | This study |
| pT7-TatABC-His | $his_6$-*tatABC* was cloned into pIVEX-2.3MCS under the control of *T7* promoter | This study |
| pT7-SufI | *sufI* was cloned into pIVEX-2.3MCS under the control of *T7* promoter | This study |
| pT7-SufI(KK) | *sufI(KK)* was cloned into pIVEX-2.3MCS under the control of *T7* promoter | This study |
| pT7-TorA-GFP | *gfp* with TorA signal sequence was cloned into pIVEX-2.3MCS under the control of *T7* promoter | This study |
| pT7-TorA-GFP(KK) | *gfp* with TorA(KK) signal sequence was cloned into pIVEX-2.3MCS under the control of *T7* promoter | This study |
| pAra-CdsA | *bla* on pKQ2-CdsA was replaced with *spc* | [4] |
| pT7-Pm-$F_0$b-His | $his_6$-$F_0b$ from *Propionigenium modestum* was cloned into pETUA under the control of *T7* promoter | This study |
| pTac-GFP | *gfp* was cloned into pUSI2 under the control of *tac* promoter | This study |

**Table S3. Oligonucleotides used in this study.**

| Name | Sequence (5'->3') | Purpose |
| --- | --- | --- |
| Tac-T(G)-His 5' in | TTTTGGATCCTAGGAGGTTTAAATTTATGAA CAATAACGATCTCT | Construction of pTac-TorA-His/pTac-TorA-GFP-His |
| Tac-TorA-His 3' in | TTTTGGTACCTTTTCAGTGATGGTGGTGATG GTGTGATTTCACCTGCGACGCGGG | Construction of pTac-TorA-His |
| Tac-SufI-His 5' in | TTTTGGATCCTAGGAGGTTTAAATTTATGTC ACTCAGTCGGCGTCAG | Construction of pTac-SufI-His |
| Tac-SufI-His 3' in | AAAGGTACCAAATTAGTGATGGTGATGGTG ATGCGGTACCGGATTGACCAACAGTTGCCC | Construction of pTac-SufI-His |
| Tac-TG-His 3' in | TTTTGGTACCTTTTTAATGGTGATGGTGATG GTGTTTGTATAGTTCATCCATGCCATGTG | Construction of pTac-TorA-GFP-His |
| Tac-T/S/G 5' ve | TCATACATAAATTTAAACCTCCTAGGATCC | Construction of pTac-TorA-His/pTac-SufI-His/pTac-TorA-GFP-His |
| Tac-TorA-His 3' ve | CCCGCGTCGCAGGTGAAATCACACCATCAC CACCATCACTGAAAAGGTACCAAAA | Construction of pTac-TorA-His |
| Tac-SufI-His 3' ve | GGGCAACTGTTGGTCAATCCGGTACCGCAT CACCATCACCATCACTAATTTGGTACCTTT | Construction of pTac-SufI-His |
| Tac-TG-His 3' ve | CATCACCATCACCATCACTAAAAAGGTACC ACCAGCGATATCCCGAACTAAAAAGGTACC | Construction of pTac-TorA-GFP-His |
| Tac-TatA-His 5' in | TTTTGGATCCTAGGAGGTTTAAATTTATGGG TGGTATCAGTATTTGGCAGTTATTGATTA | Construction of pTac-TatA-His |
| Tac-TatA-His 3' in | TTTTGGTACCTTTTTAGTGATGGTGATGGTG ATGCACCTGCTCTTTATCGTGGCGCTTCG | Construction of pTac-TatA-His |
| Tac-TatB-His 5' in | TTTTGGATCCTAGGAGGTTTAAATTTGTGTT TGATATCGGTTTTAGCGAACTGCTATTGG | Construction of pTac-TatB-His |
| Tac-TatB-His 3' in | TTTTGGTACCTTTTTAGTGATGGTGATGGTG ATGCGGTTTATCACTCGACGAAGGGGAAG | Construction of pTac-TatB-His |
| Tac-TatC-His 5' in | TTTTGGATCCTAGGAGGTTTAAATTTATGTC TGTAGAAGATACTCAACCGCTTATCACGC | Construction of pTac-TatC-His |
| Tac-TatC-His 3' in | TTTTGGTACCTTTTTAGTGATGGTGATGGTG ATGTTCTTCAGTTTTTTCGCTTTCTGCTT | Construction of pTac-TatC-His |
| Tac-TatABC 5' in | GGATCCTAGGAGGTTTAAATTTATGGGTGGT ATCAGTATTTGGCAGTTATTGATTATTGC | Construction of pTac-TatABC |
| Tac-TatABC 3' in | GGTACCTTTTTATTCTTCAGTTTTTTCGCTTT CTGCTTCAGCGTCGTTTTCCTCTTCCCG | Construction of pTac-TatABC |
| Tac-TatABC 5' ve | AATACTGATACCACCCATAAATTTAAACCTC CTAGGATCC | Construction of pTac-TatABC |
| Tac-TatABC 3' ve | AGCGAAAAAACTGAAGAATAAAAAGGTAC CACCAGCGATATCCCGAACTAAAAAGGTAC C | Construction of pTac-TatABC |
| T7-TorA-GFP 5' in | CATATGAACAATAACGATCTCTTTCAGGCAT CACGTCGGCGTTTTCTGGC | Construction of pT7-TorA-GFP/pT7-TorA-GFP-His |
| T7-TorA-GFP 3' in | CCCCCCCCGGGAGCTCGCTCGAGTCGACAA ATTATTTGTATAGTTCATCC | Construction of pT7-TorA-GFP |
| T7-TorA-GFP-His 3' in | GAACCCCCCCCGGGAGCTCGCTCGAGTCGA CAAATTAGTGATGGTGATGG | Construction of pT7-TorA-GFP-His |
| T7-TorA-GFP 5' ve | AGATCGTTATTGTTCATATGTATATCTCCTTC TTAAGTTAAACAAAATT | Construction of pT7-TorA-GFP/pT7-TorA-GFP-His |
| T7-TorA-GFP 3' ve | TAATTTGTCGACTCGAGCGAGCTCCCGGGG GGGGTTCTCATCATCATCAT | Construction of pT7-TorA-GFP/pT7-TorA-GFP-His |

| | | |
|---|---|---|
| Tac/T7-TorA-GFP(KK) 5’ | TCAGGCATCAAAAAAGCGTTTTCTGGCACA ACTCGGCGGC | Construction of pTac-TorA-GFP(KK)-His/pT7-TorA-GFP(KK) |
| Tac/T7-TorA-GFP(KK) 3’ | AACGCTTTTTTGATGCCTGAAAGAGATCGTT ATTGTTCAT | Construction of pTac-TorA-GFP(KK)-His/pT7-TorA-GFP(KK) |
| Tac/T7-SufI(KK) 5’ ve | ATGTCACTCAGTAAAAAGCAGTTCATTCAG GCATCGGGGATTG | Construction of pTac-SufI(KK)-His/pT7-SufI(KK) |
| Tac-SufI(KK) 3’ ve | CTGCTTTTTACTGAGTGACATAAATTTAAAC CTCCTAGGATCC | Construction of pTac-SufI(KK)-His |
| T7-SufI(KK) 3’ ve | CTGCTTTTTACTGAGTGACATATGTATATCTC CTTCTTAAAGT | Construction of pT7-SufI(KK) |
| T7-SufI 5’ in | TTTTAAAACATATGTCACTCAGTCGGCGTCA GTTCATTCAGGCAT | Construction of pT7-SufI |
| T7-SufI 3’ in | TTTCTCGAGAAATTACGGTACCGGATTGACC AACAGTTGCCCAAT | Construction of pT7-SufI |
| T7-SufI 5’ ve | AATGAACTGACGCCGACTGAGTGACATATG TATATCTCCTTCTTAAAGTTAAACAAAATT | Construction of pT7-SufI |
| T7-SufI 3’ ve | TTGGTCAATCCGGTACCGTAATTTCTCGAGC GAGCTCCCGGGGGGGGTTCTCAT | Construction of pT7-SufI |
| T7-TatABC 5’ in | TTTGTTTAACTTTAAGAAGGAGATATACATA TGGGTGGTATCAGTATTTGGCAGTTATTG | Construction of pT7-TatABC |
| T7-TatABC 3’ in | CCCCCCGGGAGCTCGCTCGAGTCGACTTTT TATTCTTCAGTTTTTTCGCTTTCTGCTTCA | Construction of pT7-TatABC |
| T7-TatABC 5’ ve | CAATAACTGCCAAATACTGATACCACCCATA TGTATATCTCCTTCTTAAAGTTAAACAAA | Construction of pT7-TatABC |
| T7-TatABC 3’ ve | GCAGAAAGCGAAAAAACTGAAGAATAAAA AGTCGACTCGAGCGAGCTCCCGGGGGGGG TT | Construction of pT7-TatABC |
| T7-TatABC-His 5’ | GAAGAACATCACCATCACCATCACTAATTTC TCGAGCGAGCTCCCGGGGG | Construction of pT7-TatABC-His |
| T7-TatABC-His 3’ | TTAGTGATGGTGATGGTGATGTTCTTCAGTT TTTTCGCTTTCTGCTTCAG | Construction of pT7-TatABC-His |
| T7-Pm-$F_0$b-His 5’ in | TTTGTTTAACTTTAAGAAGGAGATATACATA TGGCTCCGCAGAATATGCC | Construction of pT7-Pm-$F_0$b-His |
| T7-Pm-$F_0$b-His 3’ in | ATCGAATTCTTAATGGTGATGGTGATGGTGT TTCTCTTCCCCTACTTCGC | Construction of pT7-Pm-$F_0$b-His |
| T7-Pm-$F_0$b-His 5’ ve | CACCATCACCATCACCATTAAGAATTCGATT TCGTCGACAAGCTTAGCGG | Construction of pT7-Pm-$F_0$b-His |
| T7-Pm-$F_0$b-His 3’ ve | CATATGTATATCTCCTTCTT | Construction of pT7-Pm-$F_0$b-His |
| Tac-GFP 5’ in | AGGAAACAGGATCCTAGGAGGTTTAAATTT ATGAGCAAAGGAGAAGAACTTTTCACTGGA | Construction of pTac-GFP |
| Tac-GFP 3’ in | CCCGACTATCACCGCCCGAGATCTGTCGAC TTATTTGTATAGTTCATCCATGCCATGTGT | Construction of pTac-GFP |
| Tac-GFP 5’ ve | ACACATGGCATGGATGAACTATACAAATAAG TCGACAGATCTCGGGCGGTGATAGTCGGG | Construction of pTac-GFP |
| Tac-GFP 3’ ve | TCCAGTGAAAAGTTCTTCTCCTTTGCTCATA AATTTAAACCTCCTAGGATCCTGTTTCCT | Construction of pTac-GFP |

### Supplementary References


1. Nishiyama, K., Suzuki, T. & Tokuda, H. Inversion of the membrane topology of SecG coupled with SecA-dependent preprotein translocation. *Cell* **85**, 71–81 (1996).
2. Grant, S. G., Jessee, J., Bloom, F. R. & Hanahan, D. Differential plasmid rescue from transgenic mouse DNAs into *Escherichia coli* methylation-restriction mutants. *Proc. Natl. Acad. Sci. U S A* **87**, 4645–4649 (1990).
3. Studier, F. W. & Moffatt, B. A. Use of bacteriophage T7 RNA polymerase to direct selective high-level expression of cloned genes. *J. Mol. Biol.* **189**, 113–130 (1986).
4. Sawasato, K. *et al.* CdsA is involved in biosynthesis of glycolipid MPIase essential for membrane protein integration *in vivo*. *Sci. Rep.* **9**, 1372 (2019).
5. Nozaki, S. & Niki, H. Exonuclease III (XthA) enforces *in vivo* DNA cloning of *Escherichia coli* to create cohesive ends. *J. Bacteriol.* **201**, e00660-18 (2019).
6. Grenier, F., Matteau, D., Baby, V. & Rodrigue, S. Complete genome sequence of *Escherichia coli* BW25113. *Genome Announc.* **2**, e01038 (2014).
7. Baba, T. *et al.* Construction of *Escherichia coli* K-12 in-frame, single-gene knockout mutants: the Keio collection. *Mol. Syst. Biol.* **2**, 2006.0008 (2006).
8. Shibui, T., Uchida, M. & Teranishi, Y. A new hybrid promoter and its expression vector in *Escherichia coli*. *Agric. Biol. Chem.* **52**, 983–988 (1988).
